# Male-biased immune ageing in a polygynous bat

**DOI:** 10.64898/2026.02.27.708485

**Authors:** Jack G. Rayner, Danielle M. Adams, Najib M. El-Sayed, David M. Mosser, Gerald S. Wilkinson

**Affiliations:** Department of Biology, University of Maryland, College Park, MD 20742; Department of Cell Biology and Molecular Genetics, University of Maryland, College Park, MD 20742; Center for Bioinformatics and Computational Biology, University of Maryland, College Park, MD 20742

**Keywords:** Ageing, bats, immunity, life history, sex, social status

## Abstract

1. Bats have attracted considerable interest for their extraordinary longevity and ability to withstand infection by a range of pathogens without major harm. Yet, little is known about the relationship between these two characteristics, or the extent to which they vary between individuals of the same species.
2. We investigated sources of immune variation in wild greater spear-nosed bats, *Phyllostomus hastatus*, via transcriptome sequencing of blood samples before and after *ex vivo* exposure to lipopolysaccharide, a membrane component of gram-negative bacteria. This species exhibits an extreme harem-polygynous mating system and strongly male-biased mortality, offering the valuable opportunity to explore sources of individual variation in immunity and its relation to ageing. We assessed immune variation across both sexes, males of contrasting social status, and the full range of ages in each sex.
3. We observe striking immune variation associated with sex and age, with males and older bats mounting stronger inflammatory responses. These two factors interacted significantly, revealing male-biased slopes of age-related variation in immunity, the magnitude of which is consistent with male-biased ageing, supporting the predicted association between immunity and ageing.
4. We did not observe substantial differences in immune responses between males of harem-controlling or subordinate bachelor status. This contrasts with prior research, particularly in primates, potentially raising questions about the taxonomic generality of socially mediated immune differences, which have attracted much attention.
5. Our findings support recent calls for a nuanced approach to understanding immune adaptations and extended longevity in bats, informed by individual and species-level differences in ecology, resource allocation, and selection. These widely overlooked factors offer valuable insights into sources of immune variation and connections to other traits, such as differences in mortality and age-related deterioration.

## Introduction

Animals of the same species frequently differ markedly in their responses to immune challenges (Metcalf et al., 2025). Much of this variation likely arises because individuals differ in life history and thus resource allocation (Metcalf et al., 2020), with immune investment associated with both direct and indirect costs (Lochmiller & Deerenberg, 2000). For example, male vertebrates have been predicted to invest more in innate immune defences and inflammation than females, as their fitness is less strongly impacted by long-term physiological costs (Lee, 2006). In vertebrates, older animals are also widely expected to exhibit shifts from adaptive towards innate forms of immunity, e.g., via declining production of T cells and chronic inflammation, potentially exacerbating inflammation-driven ageing (Franceschi et al., 2000; Morrisette-Thomas et al., 2014). Yet, these individual differences are often overlooked in studies that seek to identify characteristics that underpin immune efficiency and extended longevity, which typically involve comparisons between distantly related taxa. Research that exploits variation in immune responses between individuals of the same species has the opportunity to generate insights into selection pressures and trade-offs underlying individual differences in immunity, health, and longevity at both micro- and macroevolutionary scales (Metcalf et al., 2020).

Bats have received considerable recent interest from immunologists and gerontologists (Cooper et al., 2024; Gorbunova et al., 2020; Wilkinson & South, 2002), owing to their ability to withstand infection from virulent pathogens, often without visible pathology (Munster et al., 2016; Swanepoel et al., 1996). This ability has been shaped by distinctive features of their ecology and life history. Many bats form high density colonies that facilitate pathogen transmission (Gorbunova et al., 2020; Irving et al., 2021). A common view is that this gregariousness generated selection for bats to ‘tolerate’ high frequencies of infection without mounting severe immune responses that exert harmful long-term consequences (Gorbunova et al., 2020; Pei et al., 2024), e.g., via dampened inflammasome activation (Ahn et al., 2016, 2019, 2023). Bats are also unique among mammals in being capable of powered flight, leading to elevated levels of DNA damage, which might have selected for reduced immune sensitivity to cytosolic DNA, otherwise symptomatic of viral infection (Banerjee et al., 2017; Schlee & Hartmann, 2016; Xie et al., 2018). Elevated body temperature during flight (Barclay et al., 2017; Carpenter, 1986; Reichard et al., 2010; Thomas, 1975) might also evoke a fever-like state that aids in pathogen control (O’Shea et al., 2014) or modulates antibody sensitivity (Toshkova et al., 2024), but see (Levesque et al., 2020). These immune adaptations are widely thought to be implicated in bats’ unusually long lifespans relative to their small body size (Austad, 2010), making them unusually resistant to senescence and age-related deterioration, for example, associated with harmful long-term consequences of inflammation (Huang et al., 2019). However, relevant physiological and molecular data from bats of different sexes or ages are surprisingly scarce (Cooper et al., 2024) and individual variation in immunity is rarely considered in studies of bat immune adaptations, which have tended to rely on comparative genomics (Santillán et al., 2021; Scheben et al., 2023; Zhang et al., 2013) and functional assays (Ahn et al., 2023; Banerjee et al., 2017; Xie et al., 2018) that largely ignore such variation.

We investigated variation in immune responses in wild greater spear-nosed bats, *Phyllostomus hastatus*. This species exhibits a highly polygynous mating system which is associated with strong variation in multiple life history traits, offering a valuable opportunity to assess the sources of individual differences in immune response. Males live approximately half as long as females (Adams et al., 2025), a pattern widely observed in polygynous mating systems (Staerk et al., 2025). Intense competition for mates also underlies variation in male social status, with dominant ‘harem’ males aggressively defending groups of 15 to 25 females, whereas subordinate ‘bachelor’ males congregate in more diffuse assemblages (McCracken & Bradbury, 1981), and experience high levels of intrasexual aggression (Hex and Wilkinson, in review) and accelerated mortality (Adams et al., 2025). We examined individual differences in transcriptomic responses to lipopolysaccharide (LPS): a pathogen-associated molecular pattern (PAMP) present in the cell walls of gram-negative bacteria. LPS induces potent pro-inflammatory immune responses in mammals that must be properly regulated to avoid damaging consequences, such as septic shock (Aderem & Ulevitch, 2000). We verify that exposure to LPS stimulates an inflammatory immune response, then examine differences in immune responses between sexes and males of different social statuses, which we predict to arise as a consequence of the extreme polygynous mating system, and between bats of different ages, potentially reflecting immunosenescence (Francheschi et al., 2000; Morrisette-Thomas et al., 2014). Given prior observations of male-biased rates of senescence and age-related epigenetic differences in this species (Adams et al., 2025), we predicted steeper age-related immune differences in males. Our tests of these predictions provide valuable insights into the forces that shape variation in immunity and its relationship with ageing in wild populations.

## Materials and methods

### Sampling

We collected blood from wild bats at two nearby locations in Trinidad (an abandoned ice storage building in Cumuto [10.5983°N, 61.2117°W] and a natural cave formation in Tamana [10.4711°N, 61.1958°W]), on three occasions: May 2023, January 2024, and January 2025. We captured groups of bats from depressions in cave or building ceilings using a bucket with attached laundry hamper on the end of an extendable pole. We cleaned the wing membrane of each wing with 70% isopropanol prior to collecting a 4mm diameter biopsy that was preserved in Zymo DNA/RNA Shield. Unless individual ages were known from mark-recapture records, we extracted DNA from the preserved wing biopsy for methylation profiling to estimate chronological age based on a previously published ‘all bat’ methylation clock (Wilkinson et al., 2021), which accurately predicts chronological ages of male and female *P. hastatus* (Adams et al., 2025).

To collect blood, we pierced the vein running through the wing’s propatagium using a sterile lancet and used a heparinized minivette (Sarstedt) or capillary tube (Kimble) to collect 50ul of blood. Untreated blood samples were immediately preserved by addition of 150ul of DNA/RNA Shield (Zymo), while immune-stimulated samples were treated with 5µl of lipopolysaccharide (Invivogen, cat. no. *tlrl-peklps*) for a final concentration of 1µg/mL, and incubated at 38°C for three hours before preservation of RNA by addition of 150ul of DNA/RNA Shield. Sample order was alternated to avoid confounding effects. We immediately preserved untreated samples due to possible contamination with PAMPs under non-sterile field conditions, which could confound comparisons involving untreated samples.

Animal handling methods followed guidelines by the American Society of Mammalogists and were approved by the University of Maryland Institutional Animal Care and Use Committee under licences from the Forestry Division of the Ministry of Agriculture, Land and Fisheries, Trinidad and Tobago.

### Nucleic acid extraction, library preparation and sequencing

We extracted DNA and RNA from wing punches and blood samples using Zymo Quick-DNA Miniprep and Quick-RNA Whole Blood Microprep kits, respectively, including proteinase K treatment (4 to 16 h at 60C for DNA, 1 h at room temperature for RNA). DNA samples were shipped to the Clock Foundation for methylation profiling, as described elsewhere (Adams et al., 2025; Wilkinson et al., 2021). RNA Libraries were prepared using Illumina Stranded Total RNA with Ribo-Zero Plus with 350-500ng input of high integrity (RINe > 7) total RNA and 13 PCR cycles, with globin and ribosomal RNA sequences selectively removed using custom oligo pools. Transcriptome sequencing was performed on an Illumina NextSeq 1000 platform to produce ∼15 million paired-end 58bp reads per library at the University of Maryland Brain and Behavior Institute’s Advanced Genomic Technologies Core. Adapter sequences were removed by the facility. We did not perform further trimming of reads, ∼95% of which had average quality scores > 30. Gene expression was quantified by pseudoalignment to an annotated *Phyllostomus hastatus* transcriptome (GCF_019186645.2) with Kallisto (v0.46.2) (Bray et al., 2016). We sequenced RNA from 70 untreated blood samples and 96 LPS-treated samples in 7 batches (independent sequencing runs), across 5 phases (independent batches of library preparation) (Fig. S1).

Small RNA libraries (N = 40) were prepared from RNA samples from 20 of the matched pairs of untreated and LPS-treated blood described above (10 of each sex) at the Institute for Genome Sciences, University of Maryland, Baltimore. Libraries were prepared from total RNA using Perkin Elmer Small NextFlex v4 kits and sequenced on an Illumina NovaSeq 6000 on an S4 flow cell to produce 10 million paired-end 150bp reads per library. Adapters were trimmed at the facility, with ∼99% of trimmed reads having average quality > 30. R1 reads used in analysis were further filtered using cutadapt (Martin, 2011) to retain those with length >= 18 and =< 26bp and processed using the miRDeep2 pipeline (Friedländer et al., 2012). Collapsed genome alignments were cross-referenced with miRNAs from humans, mice and three bat species (*Pteropus alecto*, *Eptesicus fuscus*, and *Artibeus jamaiciensis*) from miRbase (Kozomara & Griffiths-Jones, 2011), plus hairpin sequences of two more bat species (*Myotis myotis* and *Myotis lucifugus*) (Huang et al., 2016; Iwanowicz et al., 2013). MiRDeep2 identified 103 high-confidence putative novel miRNAs in *P. hastatus*, based on scores > 10, total read counts > 10, significant randfold P-values (indicating stable secondary structure), and no existing miRBase miRNAs with the same seed or rfam alerts. We filtered out duplicate miRNA sequences before quantifying abundance. Putative gene targets for human-annotated miRNAs were identified using TarBase v9.0 (Skoufos et al., 2024). Overrepresentation of miRNA targets was tested using default settings in miRPath v4.0 (Tastsoglou et al., 2023), after performing the differential expression analysis including only human annotated miRNAs.

### Statistical analysis

Statistical analysis was performed in R v.4.4.2. We performed differential expression/abundance analysis using DESeq2 (v1.44.0) (Love et al., 2014). To test effects of LPS treatment in each species, we ran paired analyses of blood samples from the same animal (Expression ∼ Treatment + ID): note that paired samples were sequenced in the same batch. We accounted for batch effects in all other models of *P. hastatus* gene expression via inclusion as a fixed effect. We tested for differences in gene expression associated with sex and age using models with both predictors (i.e., Expression ∼ Sex + Age + Batch), with the exception of sex-specific models. Tests of status effects in males also included age as a predictor (Expression ∼ Age + Status + Batch). Initial exploratory visualisations through variance partitioning (Hoffman & Schadt, 2016) suggested samples from different roost sites (i.e., Tamana or Cumuto) showed negligible differences in gene expression, so we did not include site in final models. We adjusted P-values for multiple testing using a Benjamini-Hochberg correction with a false discovery rate of 0.05 and did not impose fold-change criteria. We fitted further linear mixed models, using lme4 which we tested for significance using Wald’s Chi-squared statistic with type II or type III (if an interaction was included) sums of squares. Among LPS-treated samples, eleven bats were sampled on more than one occasion, whereas among untreated bat samples, three bats were sampled on more than one occasion. In each case, bats were sampled a minimum of 8 months apart. We repeated our analyses without repeated samples to check this did not influence general patterns of sex or age-related variation in immune profiles (Supplemental information).

We tested gene ontology (GO) overrepresentation of biological processes using Fisher’s exact tests against a background list of all genes included in the analysis and KEGG pathway enrichment in ClusterProfiler (Xu et al., 2024) based on human annotations (org.Hs.eg.db), due to limited functional annotation of the *P. hastatus* genome. We checked that results had similar interpretation using the P. hastatus annotation, but noted that many fewer genes were assigned to GO categories. KEGG and gene ontology overrepresentation of predicted miRNA targets was performed using miRPath 4.0 (Tastsoglou et al., 2023). We implemented weighted gene co-expression network analysis using the R package WGCNA (Langfelder & Horvath, 2008) after partitioning out the sequencing batch effect using the R package sva (Leek et al., 2012), with a minimum module size of 30 genes, a merge cut height of 0.35, and retaining 50% of the most variable genes.

### Species comparison

We retrieved publicly available RNA-sequencing data from studies that tested effects of LPS treatment on gene expression of blood samples from other mammals: leukocytes of yellow baboons (*Papio cynocephalus*) (Lea et al., 2018), and peripheral blood mononuclear cells of rhesus macaques (*Macaca mulatta*) (Snyder-Mackler et al., 2016) and pigs (*Sus domesticus*) (Li et al., 2021). The pig dataset included animals that were variable for a specific genotype, and we included only heterozygotes. Gene abundances were quantified using transcriptome references from the respective species (GCF_008728515.1, GCF_049350105.2, GCF_000003025.6, respectively) as above, with the exception of the macaque sequencing data which was available as single-end reads, so we performed single-end transcript abundance estimated in Kallisto with two sets of parameters: fragment length 180 or 200bp, and standard deviation of 20 or 30, with the true values unknown. The different parameters produced near-identical differential expression results (Pearson’s r > 0.99), so we used a fragment length of 180bp and a standard deviation of 20 in our final quantification. Genes encoding one-to-one protein orthologs were classified according to annotations from EggNog, with parameters: -m diamond, --evalue 0.001, --score 60, --pident 40, --query_cover 20, --subject_cover 20, -type proteins, –tax_scope 40674 (Mammalia), --target_orthologs one2one. Orthologs were filtered to retain only those assigned to a single annotated gene in each species.

## Results

### Immune stimulus provokes strong inflammatory responses

Treatment of blood samples with LPS provoked a strong immune response, involving core genes and pathways associated with inflammation. Principal component analysis (PCA) confirmed that LPS treatment explained substantial transcriptomic variation: untreated and LPS-treated samples were separated on PC1 (explaining 51% of variation among the top 1000 variable genes), whereas PC2 was associated with interindividual variation (Fig. 1A). LPS treatment affected expression of 7,357 genes (of 15,993 included in the analysis), with many immune-associated genes up-regulated following LPS treatment (Fig. 1B). Up-regulated genes were overrepresented for several immune-related biological processes (Fig. 1C), and enriched for key immunological and inflammatory KEGG pathways, including TNF, NF-kappa β, cytokine-cytokine receptor and IL-17 signalling (Fig. 1D). Interpretation of GO overrepresentation analysis was unchanged if we used *P. hastatus* annotations, despite many fewer genes per category (Fig. S2). Specific inflammation-associated transcription factors such as *NFKB1* (log_2_ fold-change: 2.54), and cytokines *IL1A* (log_2_FC: 7.87), *IL1B* (log_2_FC: 7.62) and *IL6* (log_2_FC: 7.59), were strongly up-regulated (P_adj._ < 1.06e-100) following LPS stimulus, in some cases contrasting with prior reports of dampened immune up-regulation in bats (Supporting information). LPS treatment affected the abundance of 59 miRNAs (of 407 included in the analysis), 30 of which were up-regulated (Fig. S3). Putative gene targets of the 29 human-annotated miRNAs that responded to LPS treatment were 3.68 times as likely to be differentially expressed in the RNA-seq data (Fisher’s test: P < 0.001) and were overrepresented for biological processes including ‘viral process’, ‘positive regulation of I-kappaB kinase/NF-kappaB signaling’ and ‘interleukin-1-mediated signaling pathway’ (all FDR< 1.94e-05); and KEGG pathways ‘shigellosis’, ‘salmonella infection’ and ‘bacterial invasion of epithelial cells’ (all FDR < 7.25e-8), consistent with expectations following immune stimulus.

**Fig. 1.**
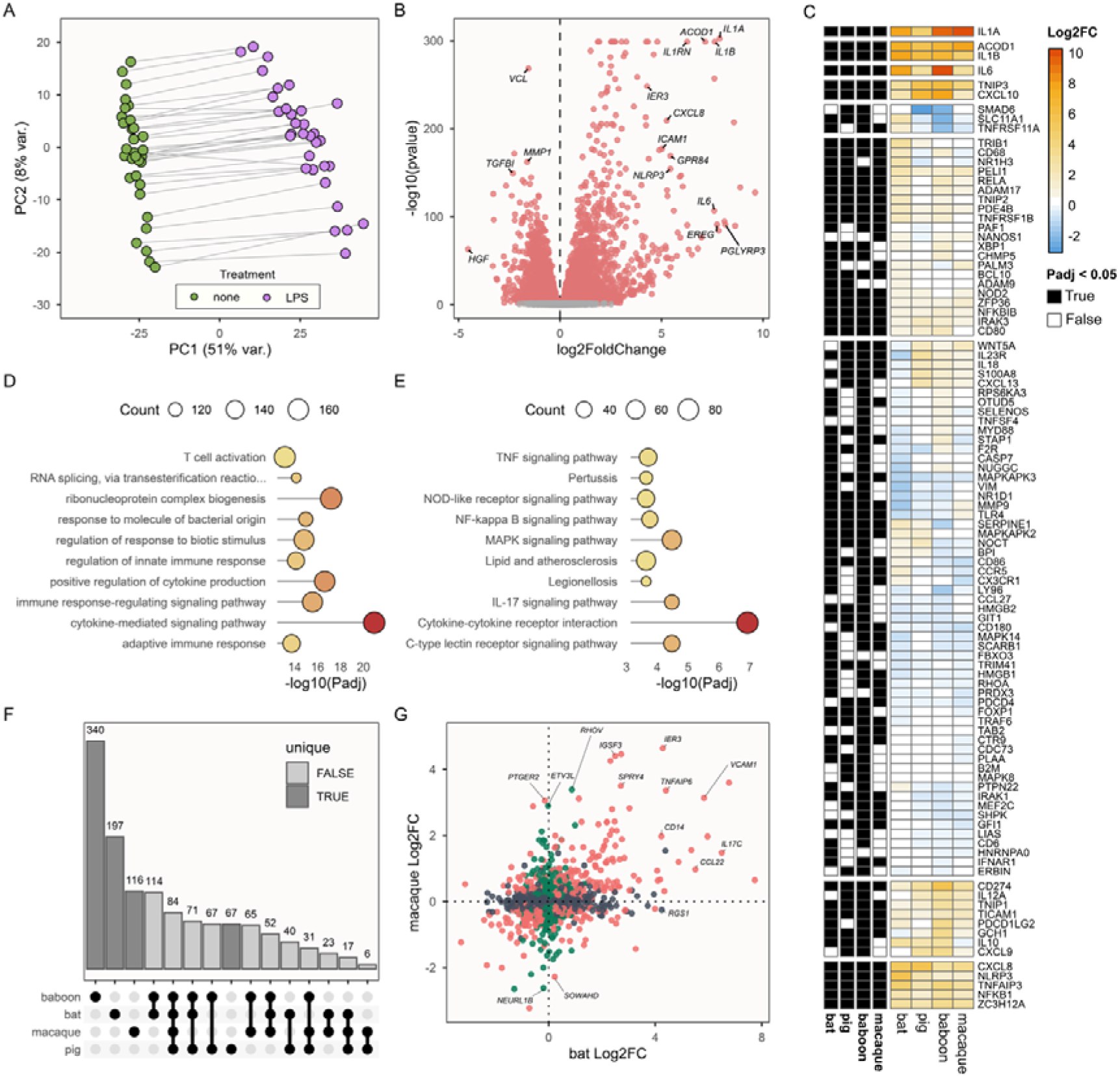
LPS affects expression of genes involved in immunity and inflammation. **A)** PCA of paired untreated and LPS-treated blood samples in bats. **B)** Volcano plot with differentially expressed genes highlighted in red. X-axis values greater than 0 indicate up-regulated expression in LPS-treated blood. **C)** The top 10 overrepresented biological process gene ontology categories, and **D)** KEGG pathway enrichment results. Point size indicates the number of DE genes in each GO category, or the number of genes in each KEGG pathway, and points are coloured according to X-axis values. **E)** Heatmap visualising log_2_ fold-changes following LPS treatment of genes in the ‘response to LPS’ gene ontology category, in four mammal species. Genes are hierarchically clustered by expression similarity. Black and white tiles indicate whether each gene was significantly differentially expressed (black) in the respective species. **F)** An ‘upset’ plot showing numbers of genes uniquely up-regulated following LPS treatment in each species (dark grey) and that are up-regulated in more than one species (light grey bars). **G)** Log_2_ fold-changes following LPS exposure in bats and macaques. Dark grey points are genes that responded significantly to LPS in bats only, green points indicate genes that responded significantly to LPS in macaques only, and peach-colored points are those that responded significantly in both species, in either direction.

To further assess whether LPS treatment stimulated the expected immune response, we compared transcriptomic responses to LPS in whole blood samples of *P. hastatus* with those of other mammals. While methodological differences (cell composition, incubation duration and LPS dose) preclude quantitative comparison, the sign and significance of responses to LPS were similar across 124 genes annotated in the ‘response to lipopolysaccharide’ gene ontology category in humans and that responded to LPS in at least one of the four species (Fig. 1E). We compared responses to LPS across 2,600 single-copy orthologous genes retained in the differential expression analysis of all four species (Fig. 1F-G). Of 598 such genes up-regulated in *P. hastatus* following LPS treatment, 401 (67%) were also up-regulated in at least one other species: 3.85 times as many as expected by chance (Fisher’s exact test: P < 0.001). These were overrepresented for a host of immunity-related biological processes (Fig. S4), whereas genes uniquely up-regulated in *P. hastatus* were not overrepresented in any category.

### Immune responses differ markedly by sex and age

We predicted that immune responses would differ between sexes and ages due to differences in life history. In support of this, large numbers of genes were differentially expressed between sexes (N = 1,826) and ages (N = 1,682) (Fig. 2A,D) across 96 LPS-treated blood transcriptomes, which is considerably more than we found for 70 untreated blood samples, in which just 132 genes were sex-biased and 28 genes exhibited age-associated variation. Substantial transcriptomic differences between sexes and ages among LPS-treated, but not untreated, blood samples therefore appear to reflect variation in immune response. Consistent with this, LPS-sensitive genes (those significantly up-regulated in blood samples following LPS-treatment) were strongly overrepresented among genes showing sex (Fisher’s exact tests; odds ratio = 3.094, P < 0.001) and age-associated (odds ratio = 1.477, P < 0.001) expression differences in immune-stimulated transcriptomes; age-associated genes. Yet, sex and age-related differences we observed in LPS-treated transcriptomes were reflected in untreated blood samples, demonstrated by significant positive Spearman rank correlations between log2 fold-changes (sex-biased genes: rho = 0.600, P < 0.001; age-associated genes: rho = 0.803, P < 0.001) (Fig. 2B,E), albeit with weaker slope (paired t-tests: P < 0.001), suggesting that immune stimulus amplified underlying differences in expression.

**Fig. 2.**
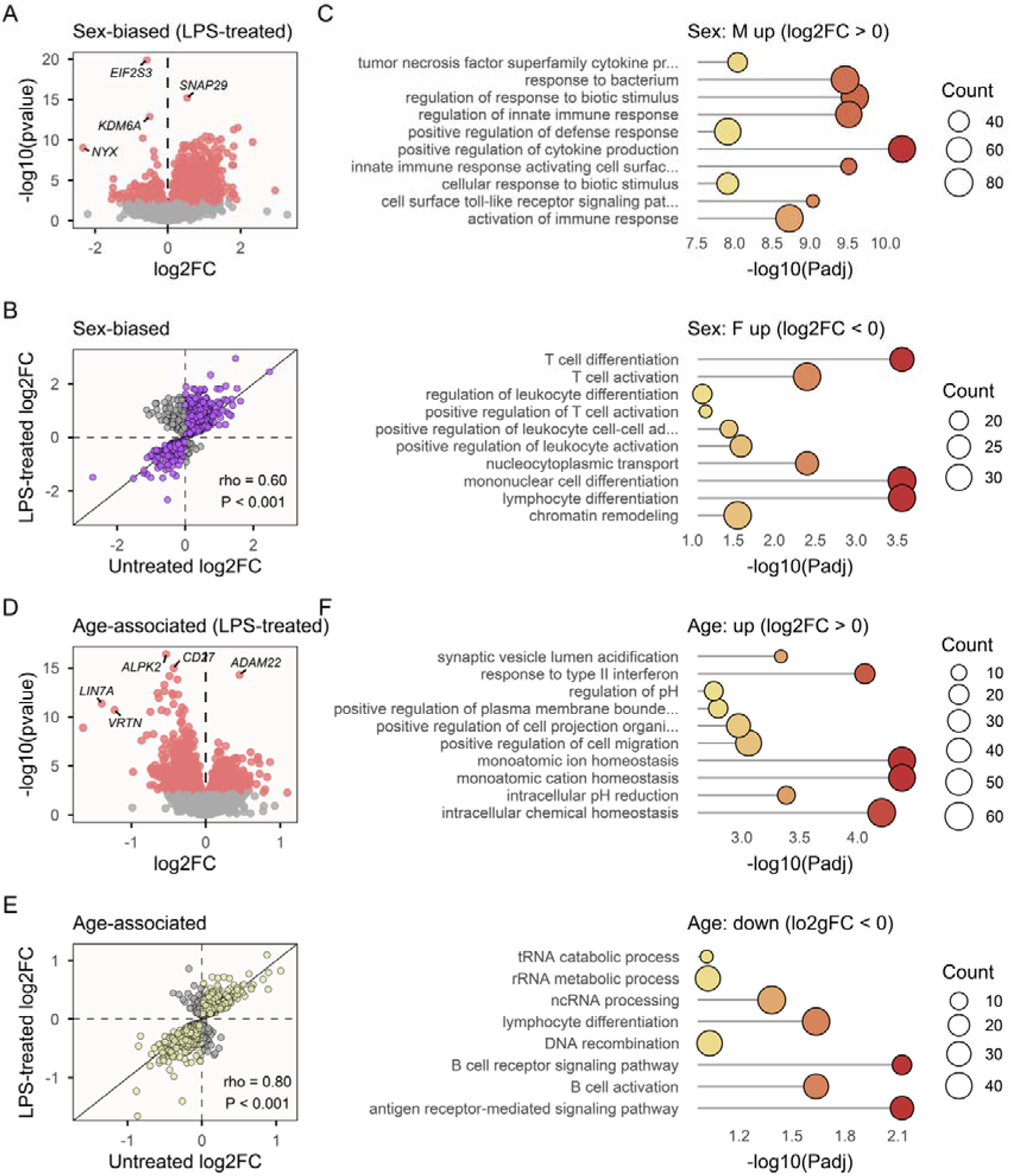
Sex and age-related differences in immune-stimulated transcriptomes. **A)** Volcano plot showing sex-differences in gene expression among LPS-treated samples, with significantly differentially expressed genes highlighted in red. **B)** Correlation between log2 fold-changes (log2FC) in untreated and LPS-treated samples across genes identified as significantly sex-biased in LPS-treated samples. Purple points are genes with the same direction of change, and Spearman rank correlations are reported across all genes. Note: three very strongly sex-biased genes (P_adj._ < 1e-86) including *Xist* and *ZFX,* that were similarly sex-biased in untreated samples, are omitted from plots A and B. **C)** Top 10 gene ontology categories overrepresented among male- and female-biased genes. **D)** Age-associated differences in gene expression among LPS-treated samples, with significantly differentially expressed genes highlighted in red. **E)** Correlation between log2 fold-changes (log2FC) in untreated and LPS-treated samples, across genes identified as significantly age-associated in LPS-treated samples. Yellow points are genes with the same sign of change, and Spearman rank correlations are reported across all genes. **F)** Top 10 gene ontology categories overrepresented among up- and down-regulated genes.

The 527 genes showing female-biased expression in immune-stimulated samples were overrepresented for genes involved in biological processes associated with adaptive immunity, such as lymphocyte differentiation and activation, whereas the 1,299 genes showing male-biased expression were over-represented for biological processes involved in innate immune responses, such as cytokine production and toll-like receptor signalling (Fig. 2C). The 771 genes that increased in expression with age were most strongly overrepresented for genes involved in ‘response to type II interferon’ and homeostasis, but were also significantly overrepresented for genes involved in ‘activation of innate immune response’ and related terms, whereas the 911 genes that declined with age were overrepresented for adaptive immunity-related processes, such as lymphocyte differentiation and antigen receptor-mediated signalling (Fig. 2F). We did not observe similar sex and age-associated variation in miRNA abundance in either untreated or LPS-treated blood (Supplemental Information).

Blood leukocyte composition influenced sex and age-related differences in immune-stimulated transcriptomes. Inclusion of log-transformed neutrophil-to-lymphocyte ratio (NLR) as a covariate in models of gene expression greatly reduced numbers of sex-biased and age-associated genes among all LPS-treated transcriptomes (from 1,826 to 49 sex-biased, and from 1,682 to 615 age-associated, genes), with NLR itself significantly associated with the expression of 5,868 genes. This suggests that gene expression profiles in LPS-treated samples are strongly influenced by differences across sexes and ages in leukocyte composition. By contrast, inclusion of NLR had little to no effect on the magnitude of differential gene expression in untreated blood samples (from 136 to 154 sex-biased, and 29 to 18 age-associated, genes), despite being associated with differential expression of 4,454 genes.

### Social status does not strongly influence male immune responses

By contrast with the substantial sex and age-related variation in immune-stimulated transcriptomes, few (N = 41) genes differed in expression between LPS-stimulated transcriptomes from 31 bachelor and 17 harem-controlling males, while accounting for differences in age, as harem-controlling males tend to be older (Wilkinson et al., 2024). These status-associated genes were not significantly overrepresented in any gene ontology categories but were enriched for genes up-regulated following LPS treatment (odds ratio = 3.485, P < 0.001). Notably, this enrichment of LPS-responsive genes was especially strong among the 22 genes that were up-regulated in harem males (odds ratio = 11.693, P < 0.001), consistent with a stronger immune response in dominant harem males, at least among this limited subset of status-associated genes. (Fig. S5)

### Males respond more strongly to immune stimulus

Male and female bats differed in their responses to LPS, but whether these differences represent immune responses of differing intensities is not immediately clear. Two initial lines of evidence supported the view that males responded more strongly to LPS. First, among paired untreated and LPS-treated samples, males tended to exhibit larger responses to LPS across LPS-sensitive genes. Sex-specific log_2_ fold-changes across these LPS-sensitive genes were strongly correlated (Pearson’s r = 0.962, df = 7355, P < 0.001; Fig. S6), but those that changed in the same direction in both sexes (the vast majority: 99.7%) exhibited a consistent male-bias in log_2_-fold change (paired t-test: t = 8.092, df = 7332, P < 0.001), with a mean difference of 0.028 (95% CI: 0.021, 0.035).

### Age-related changes in immunity occur more rapidly in males

Sex-differences in immune response might be influenced by differences in slopes of age-related variation, due to contrasting male and female longevity. To explore this possibility, we investigated sex and age-related differences across the full dataset of 70 untreated and 96 LPS-treated transcriptomes. We collapsed gene expression variation across the 1000 most variable genes using principal component analysis (Fig. 3A), which separated untreated and LPS-treated samples on PC1, and used PC1 values in LPS-treated samples as a proxy for the magnitude of response to LPS treatment (Snyder-Mackler et al., 2016). A linear mixed model incorporating sequencing batch as a random intercept showed that, by this proxy, males responded more intensely to LPS stimulus (Sex: Wald’s X^2^_1_ = 10.055, P = 0.002) and responses to LPS tended to be stronger among older bats (Age: X^2^_1_ = 3.568, P = 0.059), but did not support a sex by age interaction (X^2^_1_ = 1.449, P = 0.229; term not included in final model), though there was an indication of steeper male-biased slopes of age-associated change (Fig. 3B). We also ran a model of differences in PC1 values (i.e., *PC1_untreated_*– *PC1_LPS_*) between paired untreated and LPS-treated samples from the same blood draw (Fig. 1A), again incorporating sequencing batch as a random intercept, which did reveal a significant sex by age interaction (X^2^_1_ = 4.322, P = 0.038), with older males responding particularly strongly to LPS (Fig. 3C).

**Fig. 3.**
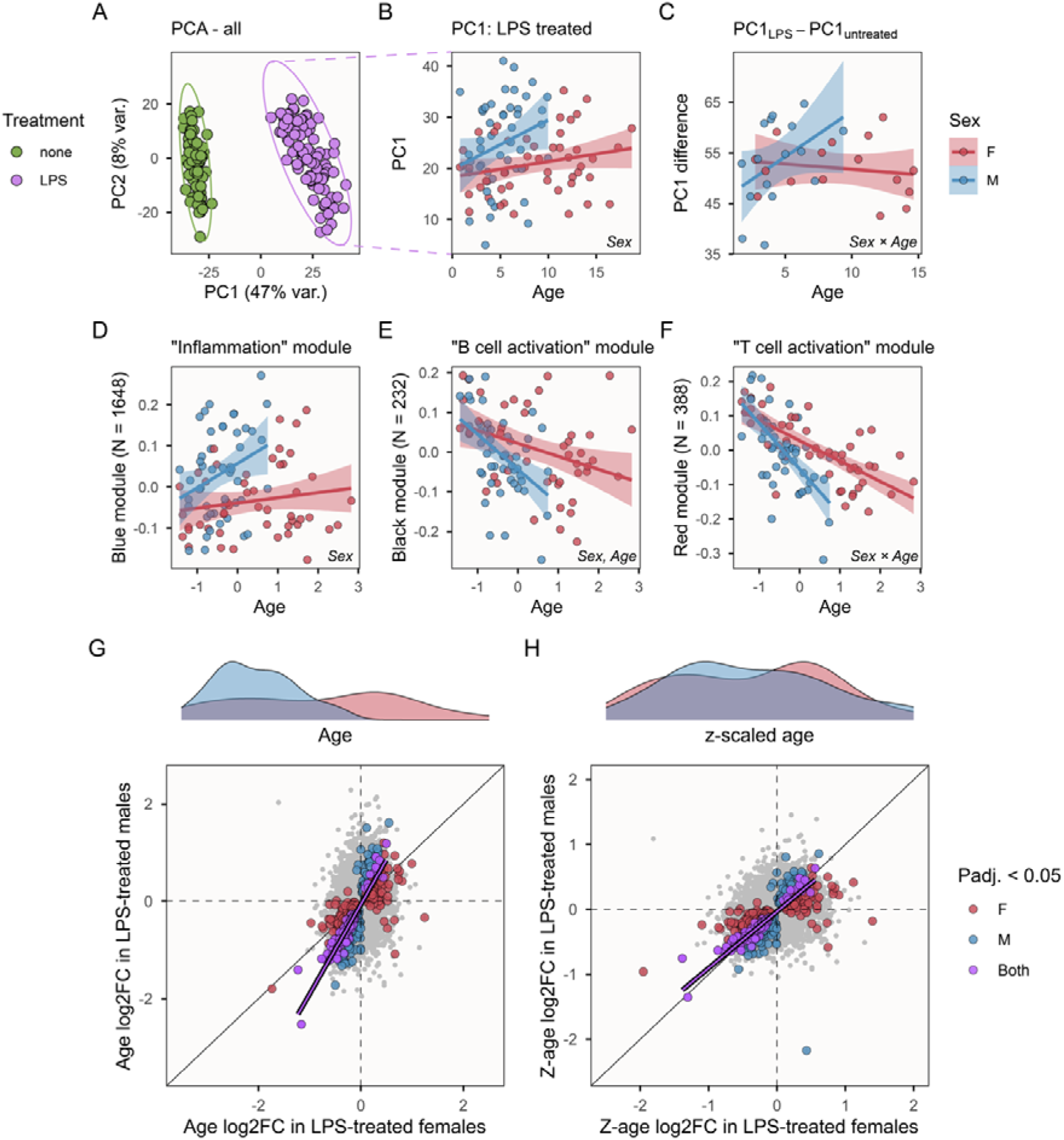
Males exhibit steeper slopes of age-related immune variation. **A)** Principal component analysis, based on the 1000 most variable genes, separates untreated and LPS-treated samples on PC1, with ellipses delineating groups of samples at 0.99 confidence level. Panel **B)** illustrates sex and age-related differences in PC1 values in LPS-treated samples, whereas panel **C)** shows differences in PC1 values between paired LPS-treated and untreated transcriptomes of the same blood sample (see Fig. 1). Panels **D-F**) show patterns of expression for representative eigengenes from gene expression modules obtained by WGCNA, labelled according to top overrepresented GO terms. Text annotations indicate significant predictor variables from the linear mixed model reported in the main text. In all panels, lines represent results of sex-specific linear regression and 95% confidence intervals. Panels **G-H)** compare slopes of significant age-related variation in gene expression between females (x-axis) and males (y-axis) in immune-treated samples, with and without sex-specific z-scaling of ages. Red and blue points show genes that are significantly age-associated females and males, respectively, with purple points significant in both sexes. The purple line shows the result of linear regression across genes that are significantly age-associated in both sexes.

To investigate the contribution of different modules of genes to sex and age-related differences in immune responses, we next implemented signed, weighted gene co-expression network analysis (WGCNA) across LPS-treated transcriptomes. This analysis identified four gene expression modules, containing between 97 and 1,648 genes, for which representative ‘eigengenes’ were associated with sex and/or age (t-test: Student’s asymptotic P_adj._ < 0.05). These included a ‘blue’ module, which was strongly overrepresented for genes involved in inflammation (Fig. S7) and was more highly expressed in males and tended to be up-regulated in older bats (Fig. 3C). Although there was no sex by age interaction (Table S1), sex-specific models reported a significant age-related expression increase in males (F_1,46_=4.613, P = 0.038), but not females (F_1,46_=1.297, P = 0.261). Further ‘black’ and ‘red’ modules were overrepresented for genes involved in B cell signalling and T cell activation/differentiation, respectively (Fig. S7). Expression of each of these modules was down-regulated in older bats and males, with a significant sex by age interaction in the red module indicating males exhibited steeper age-related decreases (Fig. 1F). The final module was more highly expressed in males and older bats, but was not significantly overrepresented for any GO biological processes. (Table S1)

Male-biased slopes of age-related variation in immune-stimulated transcriptomes based on PCA and WGCNA were reiterated by analyses at the gene level. Of 683 genes that were significantly age-associated in either sex, 665 (97.36%) showed the same direction of change in males and females (Fisher’s exact test: odds ratio = 17.18, P < 0.001). All 120 genes identified as significantly age-associated in both sexes showed concordant directions of change, and the average male slope was 1.36 ± 0.03 SE times that of females (paired t-test: t = 22.329, df = 119, P < 0.001) (Fig. 3G). Most of these consistently age-associated genes (N = 107) were negative correlated with age, and were strongly overrepresented for adaptive immunity-related biological processes (Fig. S8). Interestingly, accounting for sex-differences in age range by performing sex-specific z-scaling of ages led to the disappearance of the pattern of male-biased age-related change (t-test: P = 0.714), suggesting the immune responses of old males resemble those of old females, but with age-related changes occurring more rapidly in males (Supplemental information; Fig. 3H).

## Discussion

How bats regulate immune responses under frequent exposure to a range of pathogens has attracted considerable interest. However, a criticism recently levelled at studies of bats’ immune systems is that they often overlook species diversity in ecology and pathogen exposure (Becker et al., 2025). It is therefore not surprising that intraspecific variation has also been neglected, despite widespread appreciation of individual-level differences in immunity in animals (Habig & Archie, 2015), e.g., associated with sex (Cai et al., 2016; Kelly et al., 2018; Klein & Flanagan, 2016), age (Asquith et al., 2012; Liu et al., 2023) or social status (Lea et al., 2018; Snyder-Mackler et al., 2016). In *P. hastatus*, we observed pronounced sex and age-related differences in immunity that are consistent with our predictions: older bats down-regulated genes involved in adaptive immunity whereas older males, in particular, up-regulated genes associated with innate immunity and inflammation. This is consistent with expectations under immunosenescence (Francheschi et al., 2000; Morrisette-Thomas et al., 2014) and aligns with broader patterns of male-biased actuarial and physiological senescence in *P. hastatus*: prior research has documented sex differences in age-related variation across hormone profiles (Wilkinson et al., 2024), DNA methylation (Adams et al., 2025), telomere length (Rayner et al., 2025), and neutrophil-lymphocyte ratios (Wilkinson et al., in review). The latter is especially pertinent, as we found immune variation was strongly influenced by differential leukocyte abundance, accounting for much of the sex and age-related variation we observe in immune-stimulated samples.

Bats are widely thought to show unusual resistance to age-related deterioration, both in general and specifically with respect to immunity (Cooper et al., 2024; Gorbunova et al., 2020; Salmon et al., 2009). Our findings of considerable age-related variation in immunity in *P. hastatus* contribute to growing evidence that this is not necessarily true of all species or individuals—for instance, age-related telomere attrition, which can drive immunosenescence (Bellon & Nicot, 2017), has been observed in several bat species (Foley et al., 2018a; Forest, 2022; Ineson et al., 2020; Power et al., 2023). Our findings differ in some interesting respects from those of a study of age-related gene expression differences in blood samples of female greater mouse-eared bats (*Myotis myotis*), which reported age-related decreases in the expression of genes involved in adaptive immunity without concomitant increases in the expression of genes related to inflammation (Huang et al., 2019). The authors suggested these findings indicate bats have evolved dampened inflammatory responses that mitigate inflammation-induced ageing. By contrast, we observed transcriptomic evidence of both age-related decreases in adaptive immunity and increases in inflammation in *P. hastatus*, with the latter most pronounced in males. This difference might be influenced by species differences in ecology: *M. myotis* is a particularly long-lived bat, with a longevity quotient of 5.71, relative to 2.66 for female *P. hastatus* (1.33 in males), so it may be more resistant to age-related deterioration (Foley et al., 2018b). However, our study also likely had greater power to detect age-related differences in immunity because we assayed transcriptomic variation following immune stimulation, which greatly increased numbers of age-associated genes, and we also sampled both sexes across a wider age range, with the oldest sampled bat approximately 18 years old relative to a maximum recorded longevity of 22 years (Wilkinson & Adams, 2019). There is, nevertheless, substantial variation across bats in the severity of physiological responses to LPS (Seltmann et al., 2022), so future work incorporating multiple species is needed to illuminate the extent to which age-related changes in immunity differ or are shared across species, and the contributing factors.

As well as influencing sex differences in selection, mating systems with sex-biased reproductive skew also generate differences in social status between individuals of the same sex, which are expected to contribute to immune variation (Habig & Archie, 2015). Studies of humans and other primates have found social adversity is associated with stronger proinflammatory responses (Cole, 2014; Gassen et al., 2021; Miller et al., 2009; Snyder-Mackler et al., 2016), frequently associated with ageing phenotypes (López-Otín et al., 2023). In *P. hastatus*, bachelor and harem-controlling males differ across a range of life history traits (Adams et al., 2025; McCracken & Bradbury, 1981; Wilkinson et al., 2024), suggesting ample opportunity for social status to influence male immune profiles. Yet, we did not observe strong differences in responses to immune stimulus between these male groups. This result could challenge the taxonomic generality of the view that social status has important consequences for immunity (Campos et al., 2020), at least outside of primates and other taxa with complex social hierarchies. Even in species that do exhibit strong social differences in immunity, the nature of this relationship is variable across taxa. For example, in female macaques (*Macaca mulatta*) lower social status has been found to be associated with up-regulation of genes involved in innate immunity and inflammation (Snyder-Mackler et al., 2016), consistent with negative consequences of social adversity in humans (Snyder-Mackler et al., 2020), whereas this association is reversed in male baboons (*Papio cynocephalus*) among which higher male social status appears to be associated with proinflammatory phenotypes (Lea et al., 2018) and accelerated epigenetic ageing (Anderson et al., 2021). Additionally, lower status individuals of a variety of species display better health (Habig & Archie, 2015) and reduced parasite loads (Habig et al., 2018). In *P. hastatus*, male reproductive status is just one facet of social variation (Wilkinson, 2025). Future work also incorporating, for example, female differences in social connectivity and status could offer a more comprehensive evaluation of social differences in immunity in this species.

Recent work has emphasised that bats are not a ‘monolith’, with substantial variation across taxa in pathogen exposure and immunity (Becker et al., 2025), as well as longevity (Wilkinson & Adams, 2019) and other ecological and life history traits (Wilkinson & South, 2002). In *P. hastatus*, we observed notable differences in immune response associated with sex and age, with males exhibiting evidence of accelerated immunosenescence. To what extent this might contribute to, rather than reflect, sex-biased mortality in this species represents an important direction for future research exploring the relationship between immunity and longevity. In this and other respects, our work adds to growing calls for studies of bats and their immune systems to consider both species and individual level variation. Far from just representing statistical noise, variation between individuals of the same species provides powerful opportunities to explore factors that influence variation in immunological efficiency, with potential broader consequences for differences in ageing and longevity.

## Acknowledgements

We thank the Forestry Division of the Ministry of Agriculture, Land and Fisheries, Trinidad and Tobago for permission to capture and collect samples from greater spear-nosed bats. We thank Severine Hex, Katherine Armenta, Richard Smith, Sabira Ali, Shivam Mahadeo and Rupert Radix for assistance in the field and April Hussey for performing RNA library preparation and sequencing and for advice in planning and conducting experiments. We also thank Luke Tallon for assistance in sequencing of microRNAs and Bobby Brooke for assistance in estimating age using DNA methylation. We acknowledge the University of Maryland supercomputing resources (https://hpcc.umd.edu) and Brain and Behavior Institute sequencing facility (https://agtc.umd.edu). This work was supported by NIH grants R61-AG078474 and R33-AG078474, and NSF grant DBI-2213824.

## Data and code availability

RNA-sequencing and miRNA sequencing data are deposited at the NCBI Sequence Read Archive under the BioProject PRJNA1125049. Methylation data are available from NCBI Gene Expression Omnibus under accession GSE273259 and GSE331479. Supporting metadata and results tables are available as supporting information and alongside scripts underlying our analyses at https://github.com/jackgrayner/bat_LPS_RNA and will be made permanently available with an associated DOI via a Zenodo release upon acceptance.

## Author Contributions

GSW, DMA, DMM, NMES designed the study; GSW, DMA, JGR collected data; JGR analyzed data and wrote the initial manuscript with input from all authors; all authors contributed to writing the final manuscript.

## Supporting information

### Repeated samples

As stated in the main text, among LPS-treated samples, eleven bats were sampled on more than one occasion, whereas among untreated bat samples, three bats were sampled on more than one occasion. In each case, bats were sampled a minimum of 8 months apart, suggesting that confounding individual effects are likely to relatively minor. This number of repeated samples was insufficient to account for individual ID in our models. Instead, we repeated our analyses without repeated samples to check this did not influence general patterns of sex or age-related variation in immune profiles. Fig. S9 shows observations of sex and age-associated differences in gene expression for untreated and LPS-treated samples after repeated samples of the same bats were removed, and Fig. S10 shows the interaction of sex and age-associated variation influencing immune responses. In both cases, the interpretation is unchanged relative to that of the main text, with males and older bats up-regulating genes related to innate immunity, and with males exhibited steeper slopes of age-associated variation in immune responses.

### Responses to LPS among genes previously reported to show dampened immune responses

Interestingly, we observed clear responses to LPS treatment among genes that have previously been found to exhibit dampened immune induction in bats. For example, bone-marrow-derived macrophages and PBMCs of a pteropodid bat, *Pteropus alecto* showed dampened responses to LPS of *NLRP3*, an important component of the inflammasome, relative to humans and mice (Ahn et al., 2019). Similarly, a study of cell lines derived from big brown bats (*Eptesicus fuscus*) reported only weak up-regulation of *TNF*α, involved in systemic inflammation, following treatment with a dsRNA virus-associated PAMP, poly(I:C) (Banerjee et al., 2017). In the current study, we observed strong up-regulation of both *NLRP3* (fold-change = 43.5, P_adj._ = 2.71e-149) and *TNF*α (fold-change = 29.3, P_adj._ = 2.58e-141) in LPS-treated blood samples of *P. hastatus*. Most genes with existing annotations for involvement in LPS response showed similar responses in bats as in other mammals. A prior study of transcriptomic responses to LPS and poly(I:C) in macrophages derived from a female *M. myotis* showed that while bats up-regulate many immune-related genes as, or more aggressively than, mice, they also up-regulated the anti-inflammatory cytokine *IL-10* more strongly than other mammal species (Kacprzyk et al., 2017). However, we found that IL-10 was similarly up-regulated following LPS-treatment in *P. hastatus* as in pigs and baboons, with no significant change in macaques. It is important to note that, due to differences in methodology between studies (e.g., cell composition, incubation duration, LPS dose), we cannot directly compare magnitudes across studies. Further, LPS is a bacteria-associated molecular pattern, whereas much of the interest in bat immune defences and evidence for suppressed immune responses relates to viral infections (Gorbunova et al., 2020; Irving et al., 2021). Thus, while immune responses triggered by LPS will involve many of the same pathways as those triggered by other pathogens, our study may overlook adaptations directly implicated in dampened responses to viruses.

### Little sex or age associated variation in miRNA abundance

No miRNAs were differentially expressed between males and females in untreated or LPS-treated blood samples. With respect to age, *pal-miR-599-3p* was significantly up-regulated in LPS-treated blood samples of older bats, while *hsa-miR-7-5p* was down-regulated. In each case, age-related patterns were consistent across sexes (Fig. S11). In humans, *miR-7-5p* is involved in suppressing pro-inflammatory NF-κB activity by decreasing expression of target genes, including *IL1*β, *IL6* and *IL8* (Giles et al., 2016), so its lower abundance in older bats is consistent with reduced suppression of inflammation. Leukocyte composition did not explain a substantial proportion of variation among miRNA profiles; NLR was significantly associated with the abundance of just one miRNA, *hsa-miR-4508*, and its inclusion as a covariate did not influence patterns of sex-biased or age-associated differences.

### Sex-specific scaling of ages erodes male-biased patterns of ageing

This pattern of stronger age-related immune variation in males could be related to their elevated mortality rate, in which case old males and females might show similar immune profiles once their different lifespans are taken into account. To explore this, we reran the above analyses after performing sex-specific z-scaling of ages: in which male and female ages are each quantified in terms of standard deviations from the mean age in the respective sex (Fig. 3H cf. Fig. 3G). The interpretation of male-biased rates of immune variation subsequently dissipated: of the 120 genes identified as age-associated in the same direction in both sexes, none differed in slope after sex-specific scaling (paired t-test: t = 0.363, df = 119, P = 0.714). Similarly, sex-specific scaling of ages removed evidence of sex-by-age interactions among co-expressed gene modules.

## Supplementary figures

**Figure S1.**
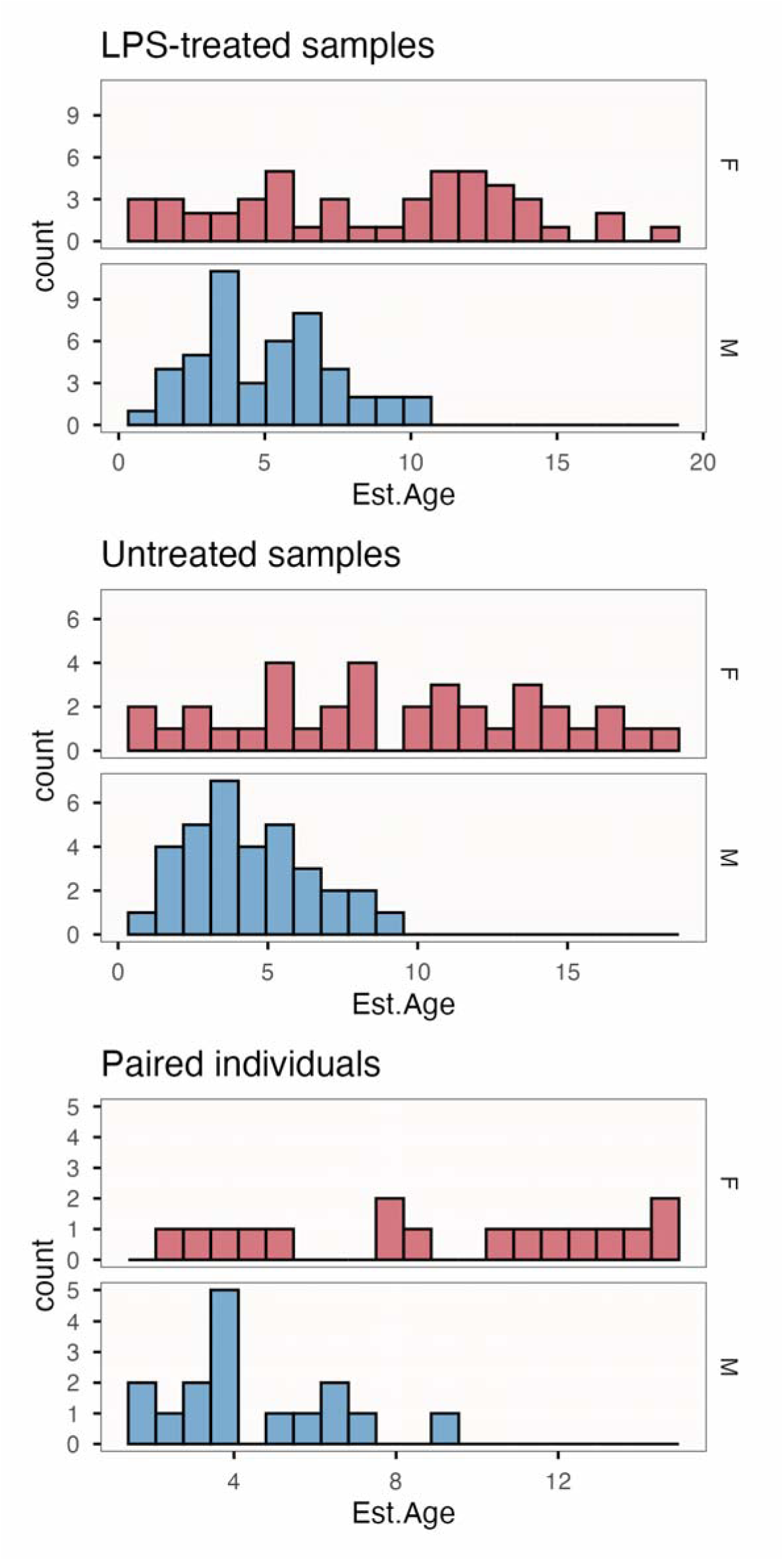
Distributions of samples across sexes and ages for untreated and LPS-treated whole-blood transcriptomes.

**Figure S2.**
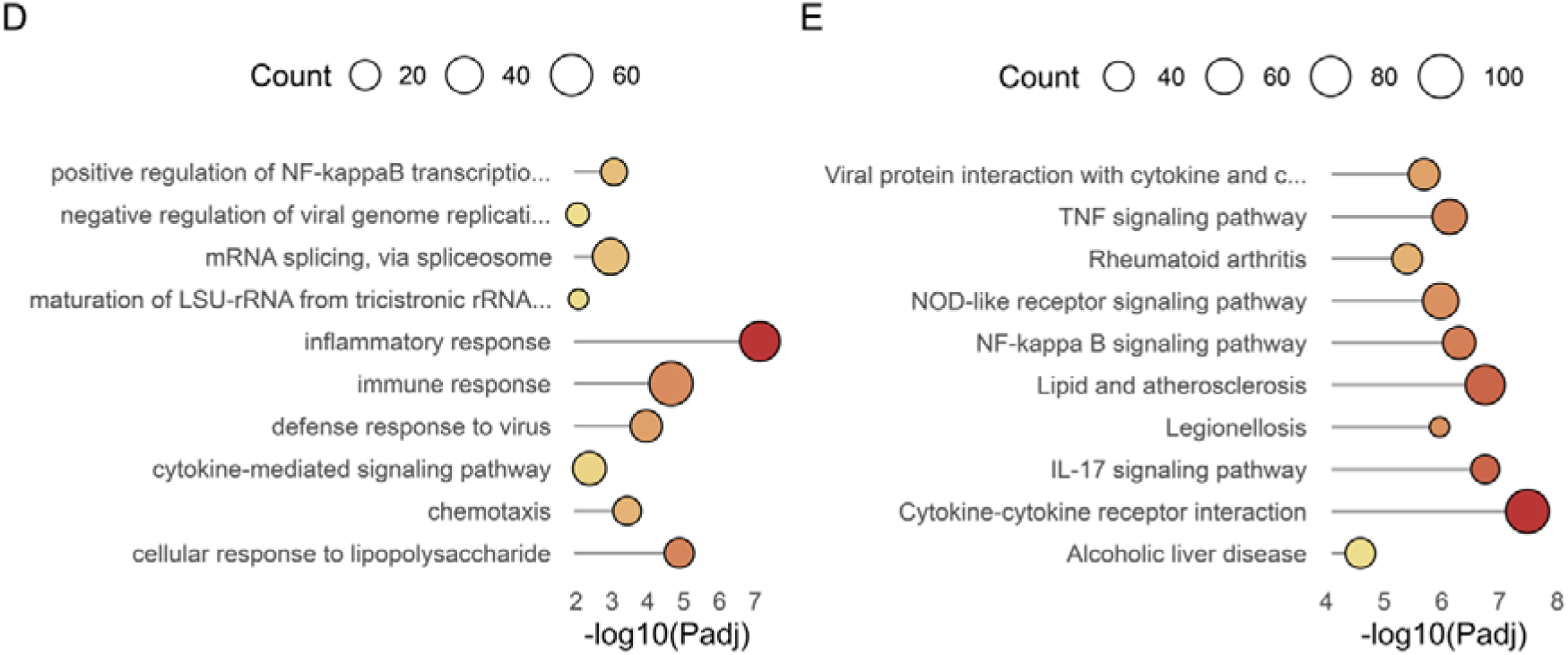
D) GO overrepresentation and E) KEGG enrichment analyses from fig. 1D,E if *P. hastatus* rather than human annotations are used. Note that, while interpretation is similar, the number of genes assigned to each GO category is considerably reduced and thus also the significance.

**Figure S3.**
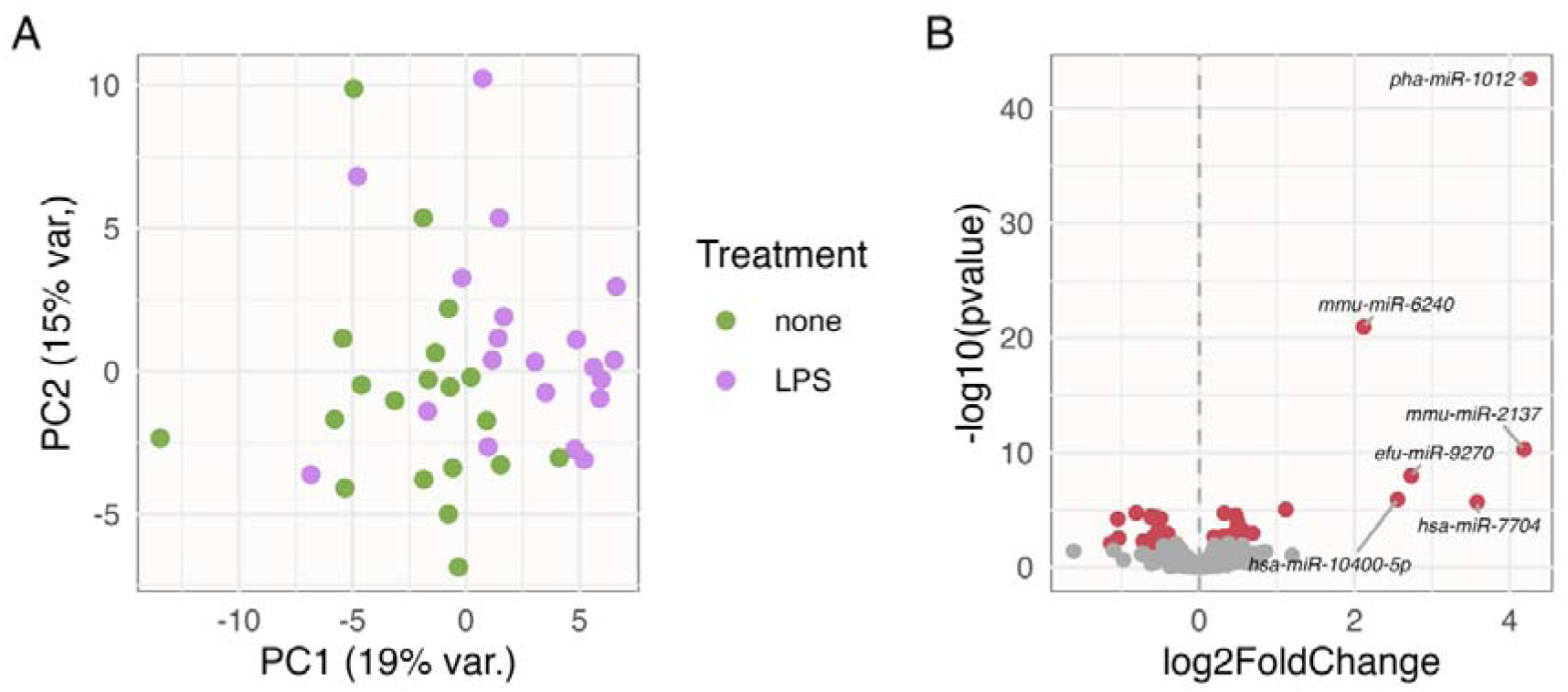
Differential expression of miRNAs following LPS-treatment. **A)** Principal component analysis across miRNA transcripts from paired untreated and LPS-treated samples. **B)** Differentially expressed miRNAs are highlighted in red.

**Figure S4.**
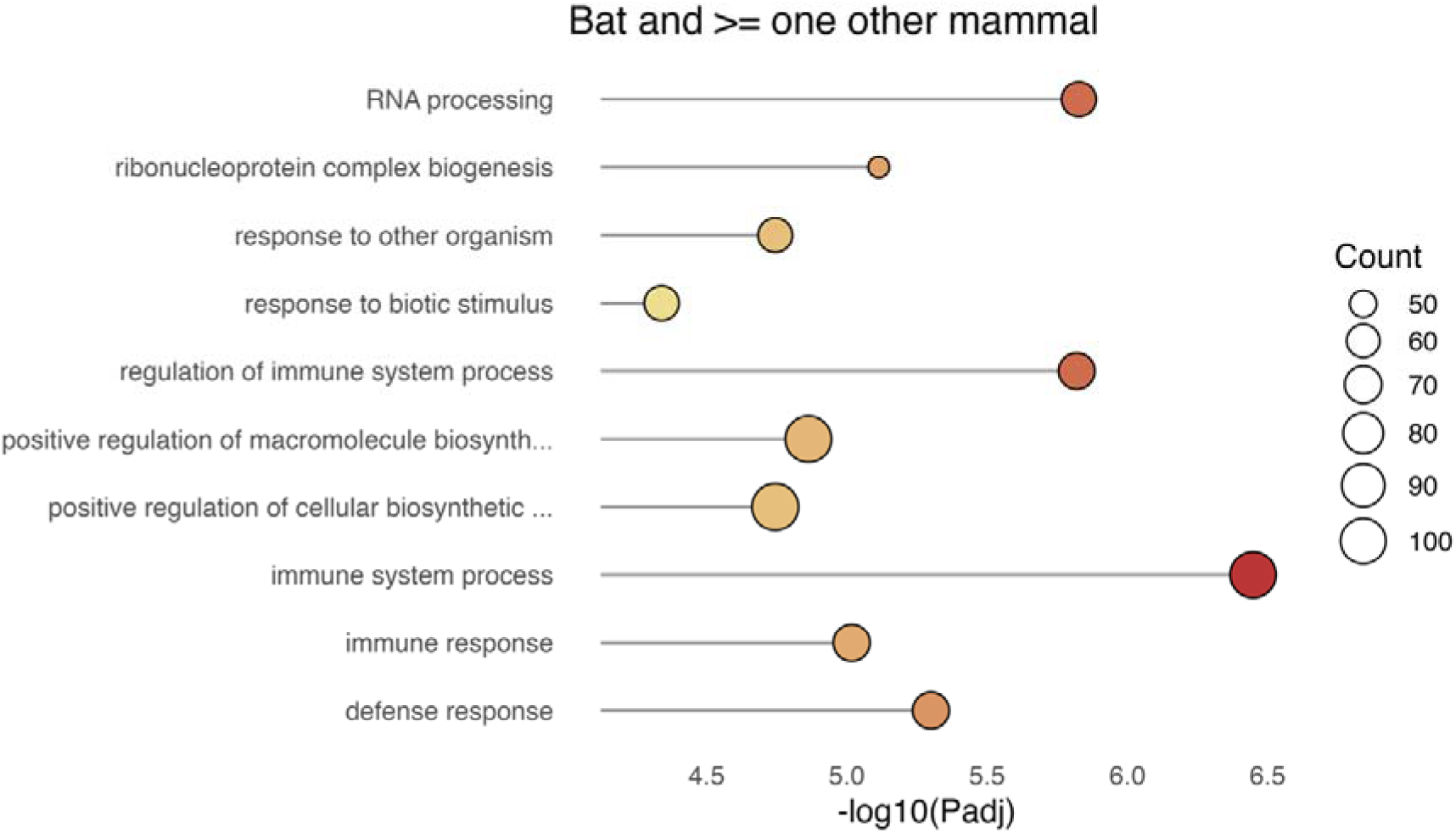
The top 10 overrepresented biological process gene ontology (GO) categories among genes that were up-regulated following LPS exposure in bats and at least one other mammal species.

**Figure S5.**
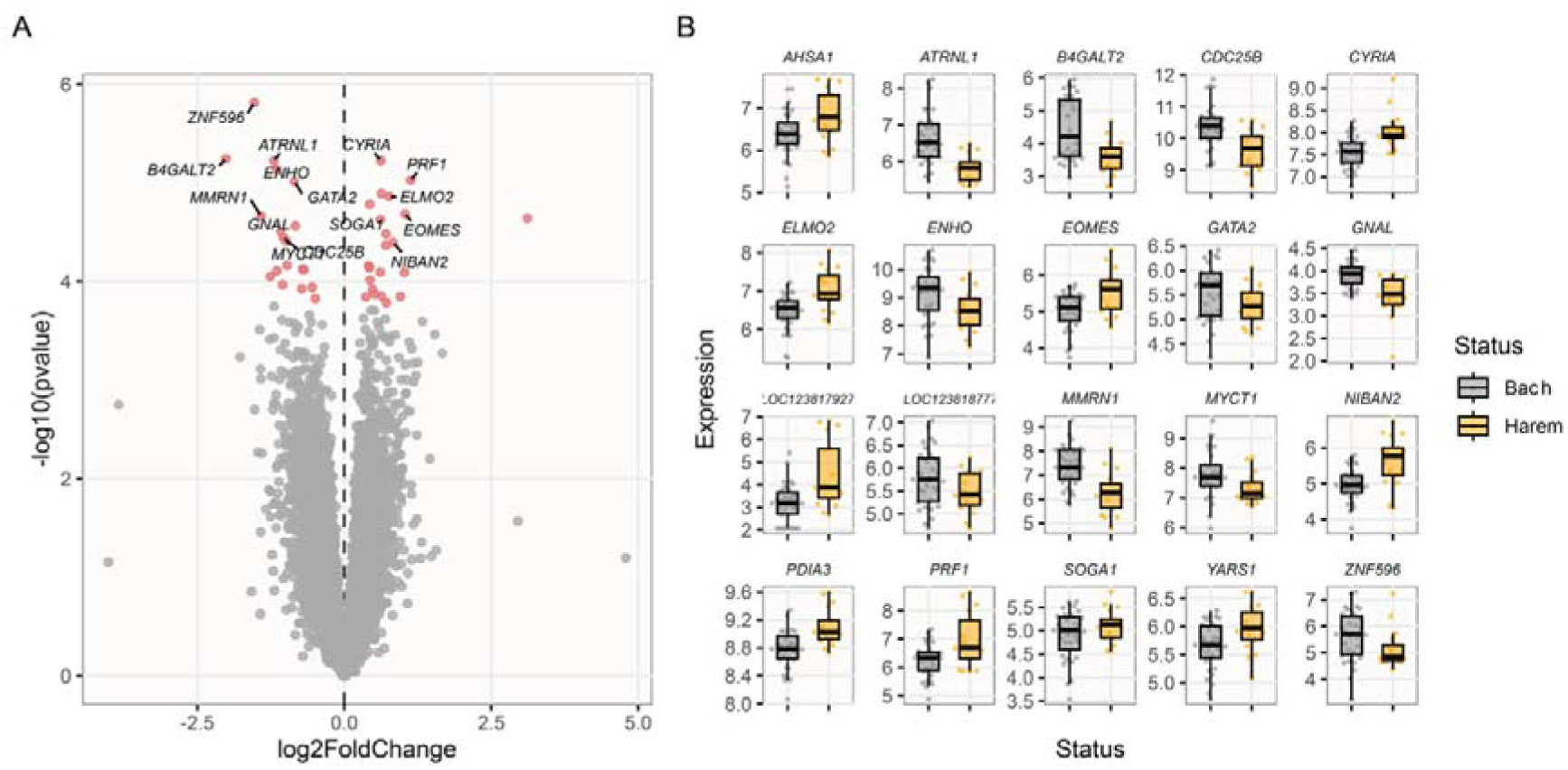
A) Volcano plot showing results of tests of differential expression between bachelor and harem males, using LPS-treated samples. Red points highlight genes differentially expressed at P_adj._ < 0.05. B) Variance stabilised expression counts across bachelor (green) and harem (yellow) males, for the top 20 differentially expressed genes.

**Figure S6.**
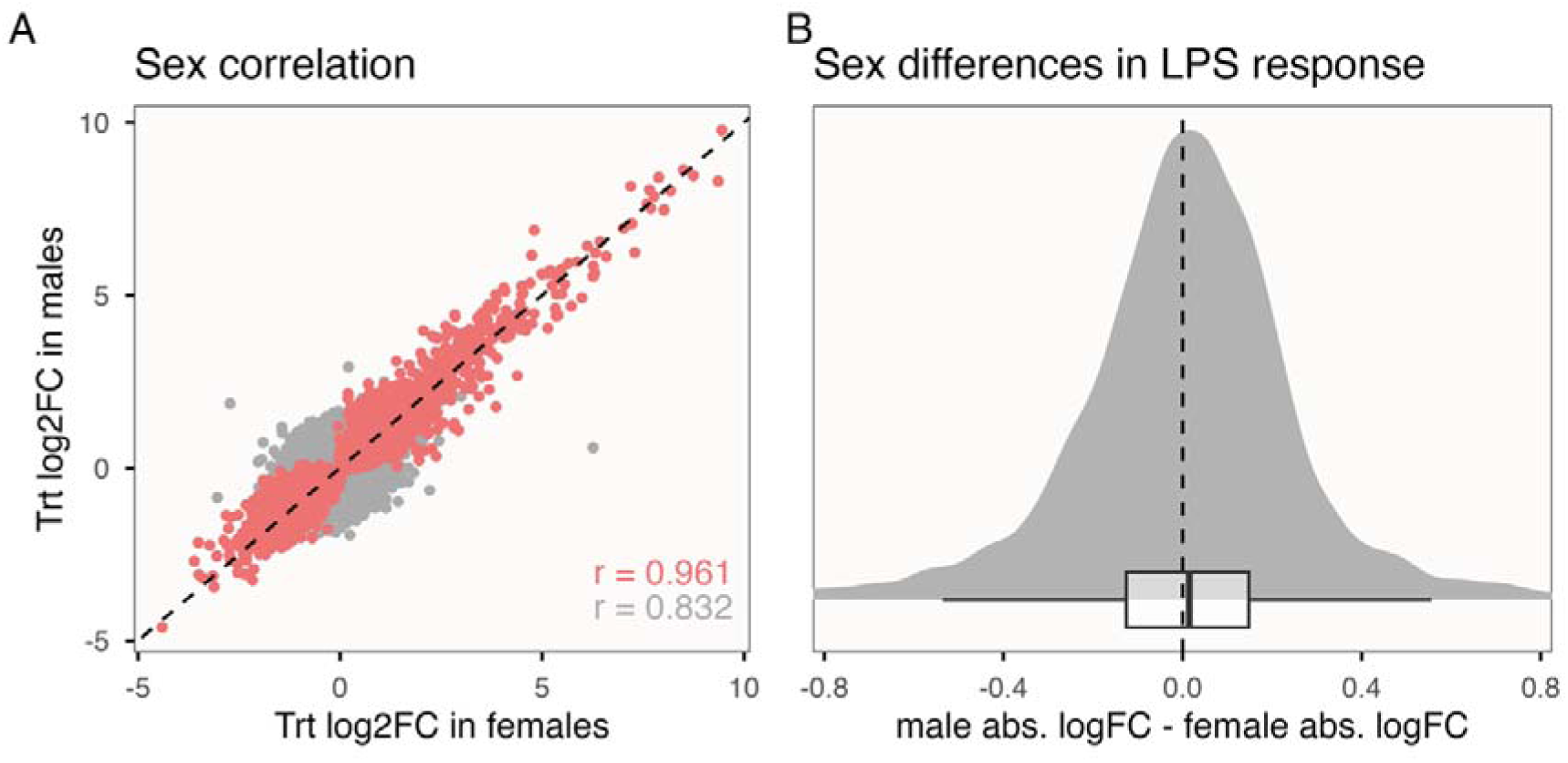
Sex-specific responses to immune stimulus. **A)** Comparison of responses to LPS treatment in females and females, with significantly LPS-affected genes highlighted in light red. Pearson’s correlation coefficients are given for significantly LPS-affected genes (red) and all genes (grey). **B)** Distribution of differences in gene-wise log_2_ fold-change across genes that responded in a consistent direction to LPS stimulus in each sex.

**Figure S7.**
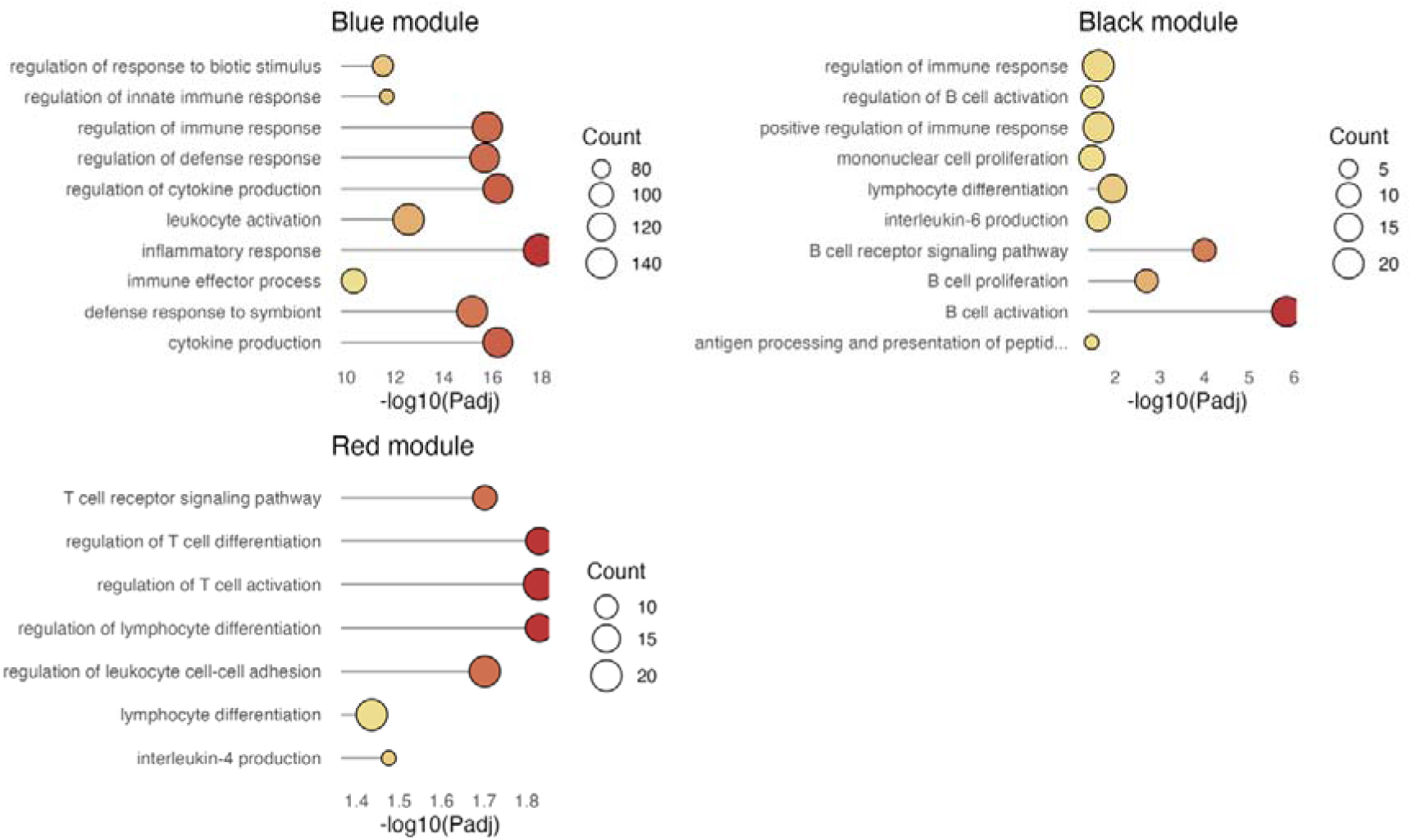
Overrepresented biological processes within each module of co-expressed genes found to be significantly associated with sex and/or age.

**Figure S8.**
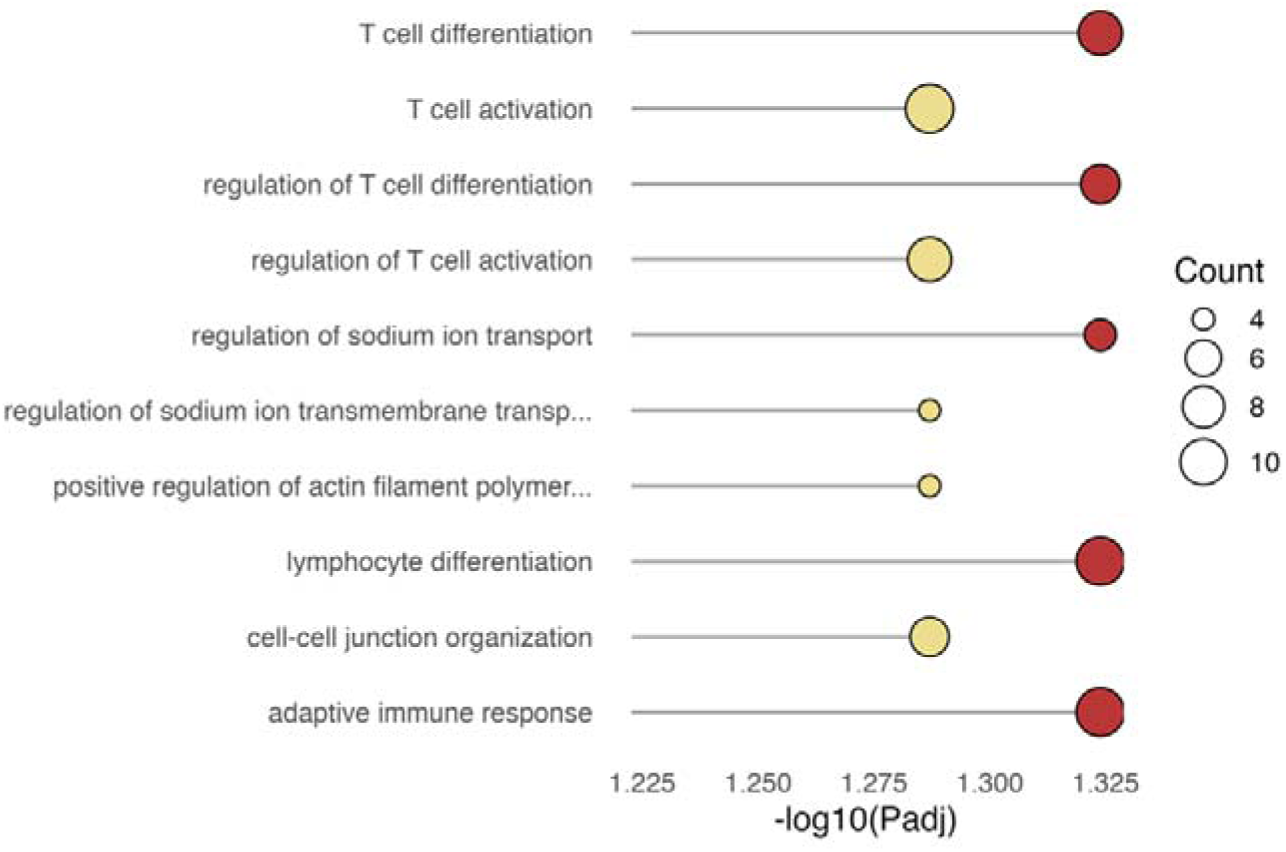
Overrepresented gene ontology categories among genes that were consistently down-regulated with age in immune-stimulated transcriptomes of both sexes.

**Figure S9.**
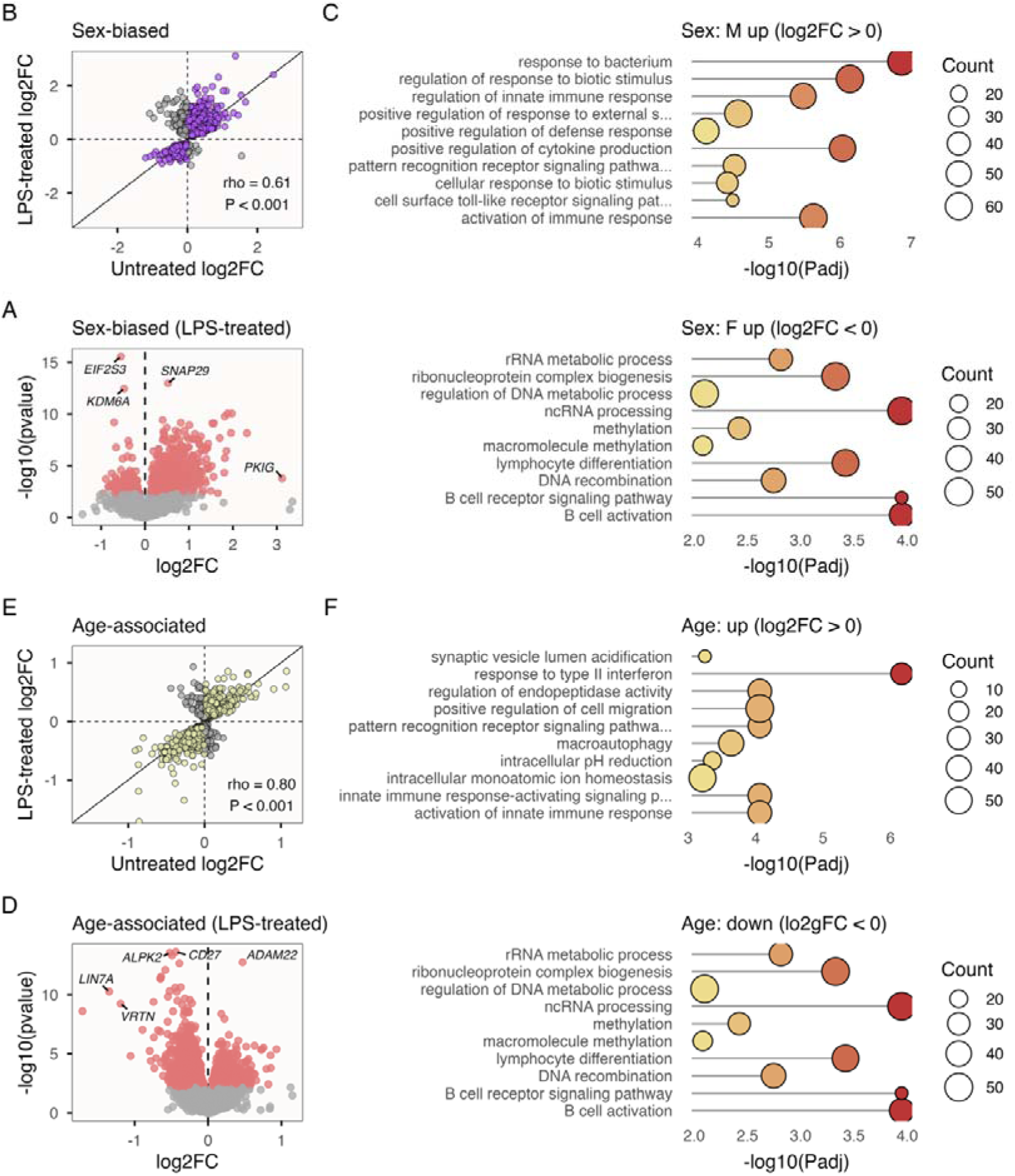
Sex and age-associated genes among LPS-treated transcriptomes, after repeated samples from the same individuals were removed.

**Figure S10.**
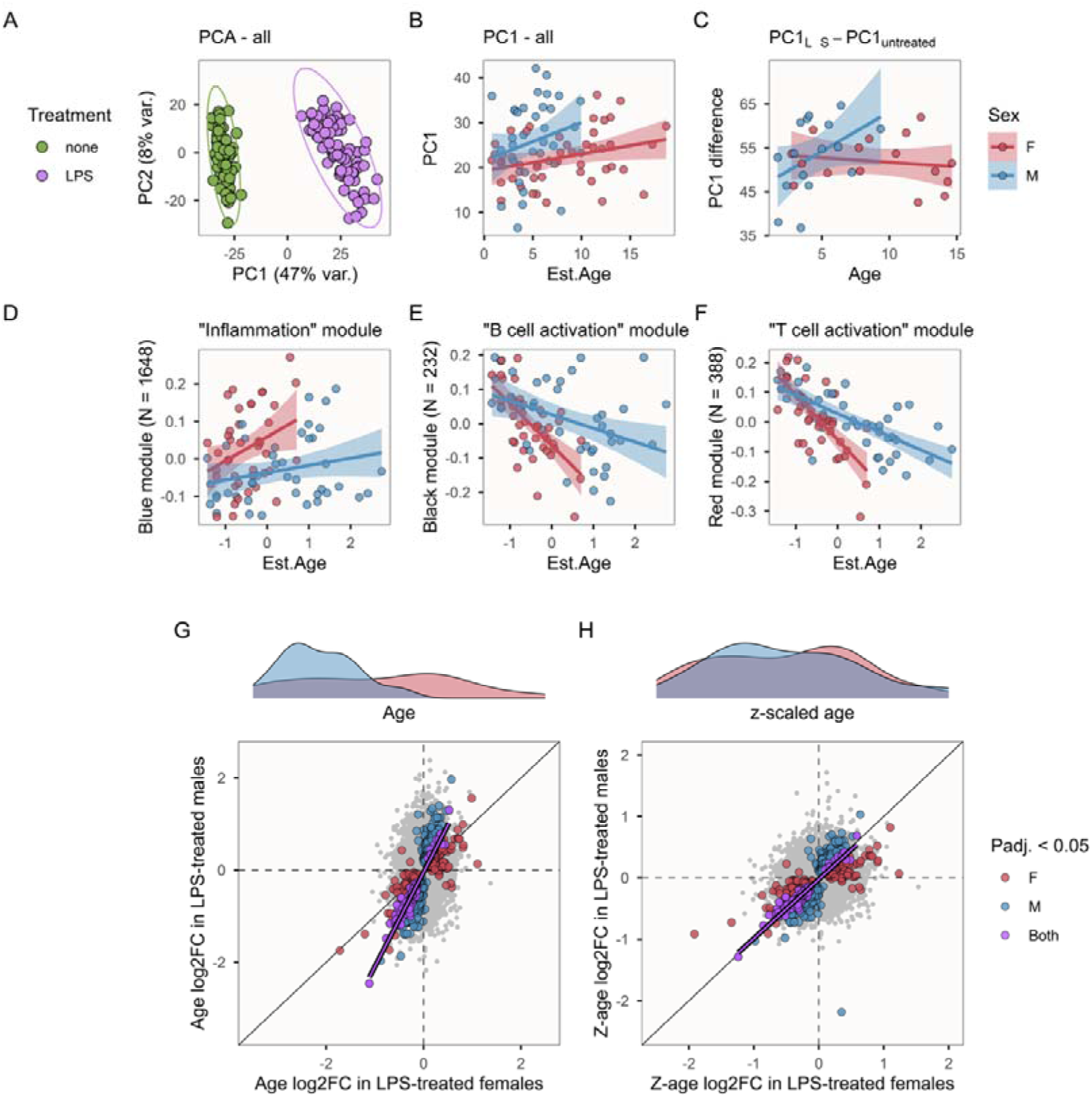
Patterns of sex and age-associated variation among LPS-treated transcriptomes (see Fig. 3), after repeated samples from the same individuals were removed.

**Figure S11.**
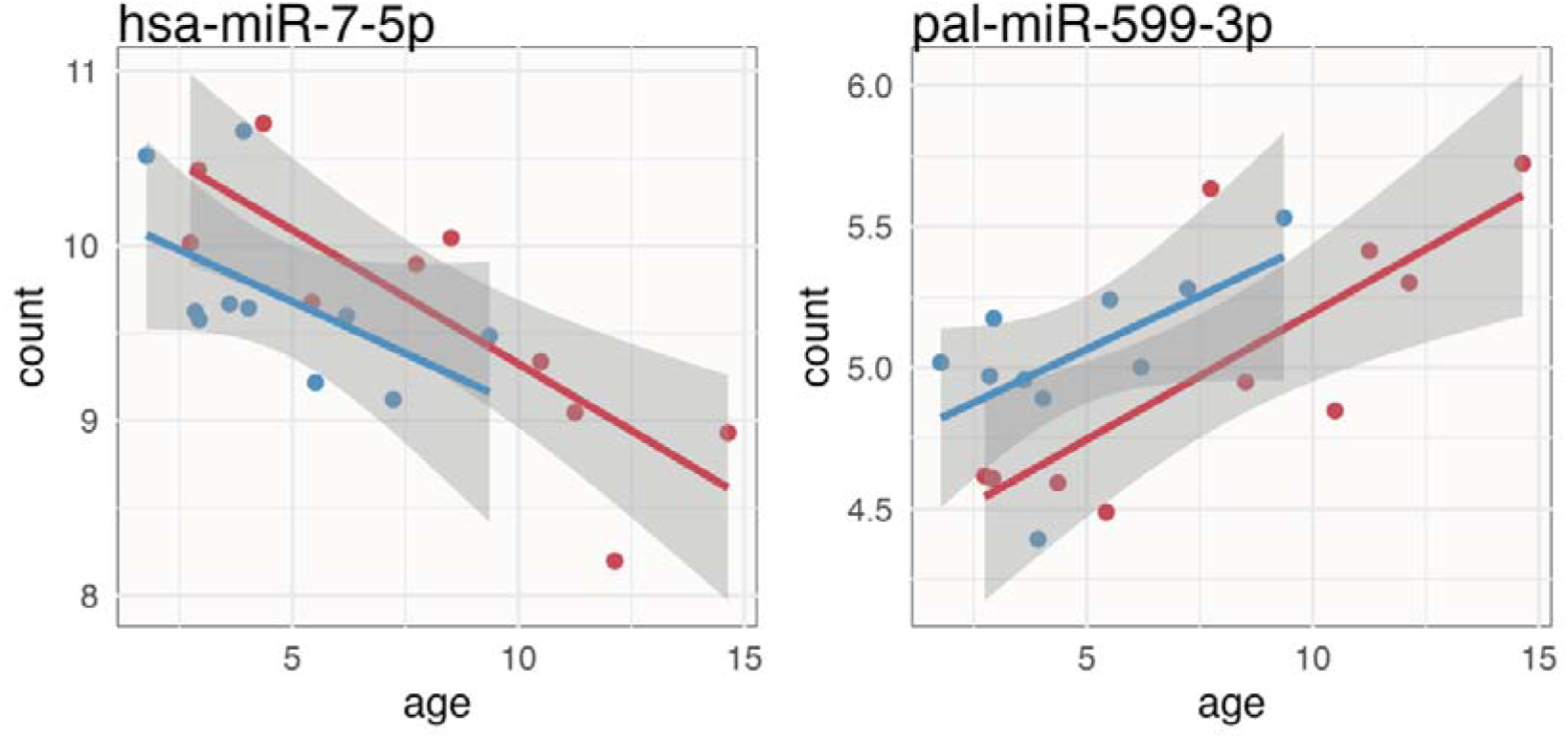
Patterns of age-associated variation in abundance of two miRNAs, showing consistent patterns in males (blue) and females (red).

## Supplementary tables

**Table S1.** Results of linear regression of WGCNA module eigengenes. Non-significant (P > 0.05) interactions were removed from models but results from interaction terms are given in footnotes. The residual degrees of freedom was 93 for models without interactions, and 92 for models with interactions.

| Module | Predictor | F value | df | P |
| --- | --- | --- | --- | --- |
| Blue<sup>1</sup> | Sex | 14.966 | 1 | < 0.001 |
|  | Age | 3.672 | 1 | 0.0584 |
| Black<sup>2</sup> | Sex | 5.587 | 1 | 0.020 |
|  | Age | 15.147 | 1 | < 0.001 |
| Red | Sex | 20.620 | 1 | < 0.001 |
|  | Age | 37.322 | 1 | < 0.001 |
| | Sex × Age | 10.587 | 1 | 0.002 |
| Magenta<sup>3</sup> | Sex | 31.638 | 1 | < 0.001 |
|  | Age | 8.430 | 1 | 0.005 |

<sup>1.</sup> Sex × Age: $F_{1,92} = 2.743$ , $P = 0.101$
<sup>2.</sup> Sex × Age: $F_{1,92} = 3.938$ , $P = 0.050$
<sup>3.</sup> Sex × Age: $F_{1,92} = 2.877$ , $P = 0.093$

## References

Adams, D. M., Rayner, J. G., Hex, S. B. S. W., & Wilkinson, G. S. (2025). DNA Methylation Dynamics Reflect Sex and Status Differences in Mortality Rates in a Polygynous Bat. *Molecular Ecology*, *n/a*(n/a), e17745. 10.1111/mec.17745

Aderem, A., & Ulevitch, R. J. (2000). Toll-like receptors in the induction of the innate immune response. Nature, 406(6797), 782–787. 10.1038/35021228

Ahn, M., Anderson, D. E., Zhang, Q., Tan, C. W., Lim, B. L., Luko, K., Wen, M., Chia, W. N., Mani, S., Wang, L. C., Ng, J. H. J., Sobota, R. M., Dutertre, C.-A., Ginhoux, F., Shi, Z.-L., Irving, A. T., & Wang, L.-F. (2019). Dampened NLRP3-mediated inflammation in bats and implications for a special viral reservoir host. Nature Microbiology, 4(5), 789–799. 10.1038/s41564-019-0371-3

Ahn, M., Chen, V. C.-W., Rozario, P., Ng, W. L., Kong, P. S., Sia, W. R., Kang, A. E. Z., Su, Q., Nguyen, L. H., Zhu, F., Chan, W. O. Y., Tan, C. W., Cheong, W. S., Hey, Y. Y., Foo, R., Guo, F., Lim, Y. T., Li, X., Chia, W. N., … Wang, L.-F. (2023). Bat ASC2 suppresses inflammasomes and ameliorates inflammatory diseases. Cell, 186(10), 2144–2159.e22. 10.1016/j.cell.2023.03.036

Ahn, M., Cui, J., Irving, A. T., & Wang, L.-F. (2016). Unique Loss of the PYHIN Gene Family in Bats Amongst Mammals: Implications for Inflammasome Sensing. Scientific Reports, 6(1), 21722. 10.1038/srep21722

Anderson, J. A., Johnston, R. A., Lea, A. J., Campos, F. A., Voyles, T. N., Akinyi, M. Y., Alberts, S. C., Archie, E. A., & Tung, J. (2021). High social status males experience accelerated epigenetic aging in wild baboons. eLife, 10, e66128. 10.7554/eLife.66128

Asquith, M., Haberthur, K., Brown, M., Engelmann, F., Murphy, A., Al-Mahdi, Z., & Messaoudi, I. (2012). Age-dependent changes in innate immune phenotype and function in rhesus macaques (Macaca mulatta). Pathobiology of Aging & Age-Related Diseases, 2(1), 18052. 10.3402/pba.v2i0.18052

Austad, S. N. (2010). Methusaleh’s Zoo: How Nature provides us with Clues for Extending Human Health Span. Proceedings of the Second Merial European Comparative Vaccinology Symposiuml: “Vaccination Challenges in Ageing Populations*,”* 142, S10–S21. 10.1016/j.jcpa.2009.10.024

Banerjee, A., Rapin, N., Bollinger, T., & Misra, V. (2017). Lack of inflammatory gene expression in bats: A unique role for a transcription repressor. Scientific Reports, 7(1), 2232. 10.1038/s41598-017-01513-w

Barclay, R. M. R., Jacobs, D. S., Harding, C. T., McKechnie, A. E., McCulloch, S. D., Markotter, W., Paweska, J., & Brigham, R. M. (2017). Thermoregulation by captive and free-ranging Egyptian rousette bats (Rousettus aegyptiacus) in South Africa. Journal of Mammalogy, 98(2), 572–578. 10.1093/jmammal/gyw234

Becker, D. J., Vicente-Santos, A., Reers, A. B., Ansil, B. R., O’Shea, M., Cummings, C. A., Roistacher, A. J., Quintela-Tizon, R. M., Pereira, M. M. T., Rosen, J., Banerjee, A., & Frank, H. K. (2025). Diverse hosts, diverse immune systems: Evolutionary variation in bat immunology. Annals of the New York Academy of Sciences, 1550(1), 151–172. 10.1111/nyas.15395

Bellon, M., & Nicot, C. (2017). Telomere Dynamics in Immune Senescence and Exhaustion Triggered by Chronic Viral Infection. Viruses, 9(10), 289. 10.3390/v9100289

Bray, N. L., Pimentel, H., Melsted, P., & Pachter, L. (2016). Near-optimal probabilistic RNA-seq quantification. Nature Biotechnology, 34(5), 525–527. 10.1038/nbt.3519

Cai, K. C., van Mil, S., Murray, E., Mallet, J.-F., Matar, C., & Ismail, N. (2016). Age and sex differences in immune response following LPS treatment in mice. *Brain*, Behavior, and Immunity, 58, 327–337. 10.1016/j.bbi.2016.08.002

Campos, F. A., Villavicencio, F., Archie, E. A., Colchero, F., & Alberts, S. C. (2020). Social bonds, social status and survival in wild baboons: A tale of two sexes. Philosophical Transactions of the Royal Society B: Biological Sciences, 375(1811), 20190621. 10.1098/rstb.2019.0621

Carpenter, R. E. (1986). Flight Physiology of Intermediate-Sized Fruit Bats (Pteropodidae). Journal of Experimental Biology, 120(1), 79–103. 10.1242/jeb.120.1.79

Cole, S. W. (2014). Human Social Genomics. PLOS Genetics, 10(8), e1004601. 10.1371/journal.pgen.1004601

Cooper, L. N., Ansari, M. Y., Capshaw, G., Galazyuk, A., Lauer, A. M., Moss, C. F., Sears, K. E., Stewart, M., Teeling, E. C., Wilkinson, G. S., Wilson, R. C., Zwaka, T. P., & Orman, R. (2024). Bats as instructive animal models for studying longevity and aging. Annals of the New York Academy of Sciences, n/a(n/a). 10.1111/nyas.15233

Foley, N. M., Hughes, G. M., Huang, Z., Clarke, M., Jebb, D., Whelan, C. V., Petit, E. J., Touzalin, F., Farcy, O., Jones, G., Ransome, R. D., Kacprzyk, J., O’Connell, M. J., Kerth, G., Rebelo, H., Rodrigues, L., Puechmaille, S. J., & Teeling, E. C. (2018a). Growing old, yet staying young: The role of telomeres in bats’ exceptional longevity. Science Advances, 4(2). 10.1126/sciadv.aao0926

Foley, N. M., Hughes, G. M., Huang, Z., Clarke, M., Jebb, D., Whelan, C. V., Petit, E. J., Touzalin, F., Farcy, O., Jones, G., Ransome, R. D., Kacprzyk, J., O’Connell, M. J., Kerth, G., Rebelo, H., Rodrigues, L., Puechmaille, S. J., & Teeling, E. C. (2018b). Growing old, yet staying young: The role of telomeres in bats’ exceptional longevity. Science Advances, 4(2), Article 2. 10.1126/sciadv.aao0926

Forest, E. E. (2022). Relationships Among Age, Relative Telomere Length, and DNA Methylation in Pteropus Bats. Grand Valley State University.

Franceschi, C., Bonafè, M., Valensin, S., Olivieri, F., De Luca, M., Ottaviani, E., & De Benedictis, G. (2000). Inflamm-aging: An Evolutionary Perspective on Immunosenescence. Annals of the New York Academy of Sciences, 908(1), 244– 254. 10.1111/j.1749-6632.2000.tb06651.x

Francheschi, C., Bonafe, M., Valensin, S., Olivieri, F., De Luca, M., Ottaviani, E., & De Benedictis, G. (2000). Inflamm-aging: An Evolutionary Perspective on Immunosenescence. Annals of the New York Academy of Sciences, 908(1), 244– 254. 10.1111/j.1749-6632.2000.tb06651.x

Friedländer, M. R., Mackowiak, S. D., Li, N., Chen, W., & Rajewsky, N. (2012). miRDeep2 accurately identifies known and hundreds of novel microRNA genes in seven animal clades. Nucleic Acids Research, 40(1), 37–52. 10.1093/nar/gkr688

Gassen, J., White, J. D., Peterman, J. L., Mengelkoch, S., Proffitt Leyva, R. P., Prokosch, M. L., Eimerbrink, M. J., Brice, K., Cheek, D. J., Boehm, G. W., & Hill, S. E. (2021). Sex differences in the impact of childhood socioeconomic status on immune function. Scientific Reports, 11(1), 9827. 10.1038/s41598-021-89413-y

Gorbunova, V., Seluanov, A., & Kennedy, B. K. (2020). The World Goes Bats: Living Longer and Tolerating Viruses. Cell Metabolism, 32(1), Article 1. 10.1016/j.cmet.2020.06.013

Habig, B., & Archie, E. A. (2015). Social status, immune response and parasitism in males: A meta-analysis. Philosophical Transactions of the Royal Society B: Biological Sciences, 370(1669), 20140109. 10.1098/rstb.2014.0109

Habig, B., Doellman, M. M., Woods, K., Olansen, J., & Archie, E. A. (2018). Social status and parasitism in male and female vertebrates: A meta-analysis. Scientific Reports, 8(1), 3629. 10.1038/s41598-018-21994-7

Hoffman, G. E., & Schadt, E. E. (2016). variancePartition: Interpreting drivers of variation in complex gene expression studies. BMC Bioinformatics, 17(1), 483. 10.1186/s12859-016-1323-z

Huang, Z., Jebb, D., & Teeling, E. C. (2016). Blood miRNomes and transcriptomes reveal novel longevity mechanisms in the long-lived bat, Myotis myotis. BMC Genomics, 17(1), 906. 10.1186/s12864-016-3227-8

Huang, Z., Whelan, C. V., Foley, N. M., Jebb, D., Touzalin, F., Petit, E. J., Puechmaille, S. J., & Teeling, E. C. (2019). Longitudinal comparative transcriptomics reveals unique mechanisms underlying extended healthspan in bats. Nature Ecology and Evolution, 3(7), Article 7. 10.1038/s41559-019-0913-3

Ineson, K. M., O’Shea, T. J., Kilpatrick, C. W., Parise, K. L., & Foster, J. T. (2020). Ambiguities in using telomere length for age determination in two North American bat species. Journal of Mammalogy, 101(4), Article 4. 10.1093/jmammal/gyaa064

Irving, A. T., Ahn, M., Goh, G., Anderson, D. E., & Wang, L.-F. (2021). Lessons from the host defences of bats, a unique viral reservoir. Nature, 589(7842), 363–370. 10.1038/s41586-020-03128-0

Iwanowicz, D. D., Iwanowicz, L. R., Hitt, N. P., & King, T. L. (2013). Differential expression profiles of microRNA in the little brown bat (Myotis lucifugus) associated with white nose syndrome affected and unaffected individuals (Report Nos. 2013–1099; Open-File Report, p. 20). USGS Publications Warehouse. 10.3133/ofr20131099

Kelly, C. D., Stoehr, A. M., Nunn, C., Smyth, K. N., & Prokop, Z. M. (2018). Sexual dimorphism in immunity across animals: A meta-analysis. Ecology Letters, 21(12), 1885–1894. 10.1111/ele.13164

Klein, S. L., & Flanagan, K. L. (2016). Sex differences in immune responses. Nature Reviews Immunology, 16(10), 626–638. 10.1038/nri.2016.90

Kozomara, A., & Griffiths-Jones, S. (2011). miRBase: Integrating microRNA annotation and deep-sequencing data. Nucleic Acids Research, 39(suppl_1), D152–D157. 10.1093/nar/gkq1027

Langfelder, P., & Horvath, S. (2008). WGCNA: an R package for weighted correlation network analysis. BMC Bioinformatics, 9(1), 559. 10.1186/1471-2105-9-559

Lea, A. J., Akinyi, M. Y., Nyakundi, R., Mareri, P., Nyundo, F., Kariuki, T., Alberts, S. C., Archie, E. A., & Tung, J. (2018). Dominance rank-associated gene expression is widespread, sex-specific, and a precursor to high social status in wild male baboons. Proceedings of the National Academy of Sciences, 115(52), E12163–E12171. 10.1073/pnas.1811967115

Lee, K. A. (2006). Linking immune defenses and life history at the levels of the individual and the species. Integrative and Comparative Biology, 46(6), 1000–1015. 10.1093/icb/icl049

Leek, J. T., Johnson, W. E., Parker, H. S., Jaffe, A. E., & Storey, J. D. (2012). The sva package for removing batch effects and other unwanted variation in high-throughput experiments. Bioinformatics, 28(6), 882–883. 10.1093/bioinformatics/bts034

Levesque, D. L., Boyles, J. G., Downs, C. J., & Breit, A. M. (2020). High Body Temperature is an Unlikely Cause of High Viral Tolerance in Bats. Journal of Wildlife Diseases, 57(1), 238–241. 10.7589/JWD-D-20-00079

Li, Z., Trakooljul, N., Hadlich, F., Ponsuksili, S., Wimmers, K., & Murani, E. (2021). Transcriptome analysis of porcine PBMCs reveals lipopolysaccharide-induced immunomodulatory responses and crosstalk of immune and glucocorticoid receptor signaling. Virulence, 12(1), 1808–1824. 10.1080/21505594.2021.1948276

Liu, Z., Liang, Q., Ren, Y., Guo, C., Ge, X., Wang, L., Cheng, Q., Luo, P., Zhang, Y., & Han, X. (2023). Immunosenescence: Molecular mechanisms and diseases. Signal Transduction and Targeted Therapy, 8(1), 200. 10.1038/s41392-023-01451-2

Lochmiller, R. L., & Deerenberg, C. (2000). Trade-offs in evolutionary immunology: Just what is the cost of immunity? Oikos, 88(1), 87–98. 10.1034/j.1600-0706.2000.880110.x

López-Otín, C., Blasco, M. A., Partridge, L., Serrano, M., & Kroemer, G. (2023). Hallmarks of aging: An expanding universe. Cell, 186(2), 243–278. 10.1016/j.cell.2022.11.001

Love, M. I., Huber, W., & Anders, S. (2014). Moderated estimation of fold change and dispersion for RNA-seq data with DESeq2. Genome Biology. 10.1186/s13059-014-0550-8

Martin, M. (2011). Cutadapt removes adapter sequences from high-throughput sequencing reads. EMBnet.Journal, 17(1), Article 1. 10.14806/ej.17.1.200

McCracken, G. F., & Bradbury, J. W. (1981). Social Organization and Kinship in the Polygynous Bat Phyllostomus hastatus. Behavioral Ecology and Sociobiology, 8(1), Article 1.

Metcalf, C. J. E., Graham, A. L., Yates, A. J., & Cummings, D. A. T. (2025). Convergence and divergence of individual immune responses over the life course. Science, 389(6760), 604–609. 10.1126/science.ady9543

Metcalf, C. J. E., Roth, O., & Graham, A. L. (2020). Why leveraging sex differences in immune trade-offs may illuminate the evolution of senescence. Functional Ecology, 34(1), 129–140. 10.1111/1365-2435.13458

Miller, G. E., Chen, E., Fok, A. K., Walker, H., Lim, A., Nicholls, E. F., Cole, S., & Kobor, M. S. (2009). Low early-life social class leaves a biological residue manifested by decreased glucocorticoid and increased proinflammatory signaling. Proceedings of the National Academy of Sciences, 106(34), 14716–14721. 10.1073/pnas.0902971106

Morrisette-Thomas, V., Cohen, A. A., Fülöp, T., Riesco, É., Legault, V., Li, Q., Milot, E., Dusseault-Bélanger, F., & Ferrucci, L. (2014). Inflamm-aging does not simply reflect increases in pro-inflammatory markers. Mechanisms of Ageing and Development, 139, 49–57. 10.1016/j.mad.2014.06.005

Munster, V. J., Adney, D. R., van Doremalen, N., Brown, V. R., Miazgowicz, K. L., Milne-Price, S., Bushmaker, T., Rosenke, R., Scott, D., Hawkinson, A., de Wit, E., Schountz, T., & Bowen, R. A. (2016). Replication and shedding of MERS-CoV in Jamaican fruit bats (Artibeus jamaicensis). Scientific Reports, 6(1), 21878. 10.1038/srep21878

O’Shea, T., Cryan, P., Cunningham, A., Fooks, A., Hayman, D. T. S., Luis, A., Peel, A., Plowright, R., & Wood, J. L. N. (2014). Bat Flight and Zoonotic Viruses. Emerging Infectious Disease Journal, 20(5), 741. 10.3201/eid2005.130539

Pei, G., Balkema-Buschmann, A., & Dorhoi, A. (2024). Disease tolerance as immune defense strategy in bats: One size fits all? PLOS Pathogens, 20(9), e1012471. 10.1371/journal.ppat.1012471

Power, M. L., D, R. R., Sébastien, R., Luke, R., Gareth, J., & Teeling, E. C. (2023). Hibernation telomere dynamics in a shifting climate: Insights from wild greater horseshoe bats. Proceedings of the Royal Society B: Biological Sciences, 290, 20231589.

Rayner, J. G., Marshall, A., Adams, D. M., Kaiser, J., Armenta, K., & Wilkinson, G. S. (2025). Sex Differences in Telomere Length in a Bat With Female-Biased Longevity. Ecology and Evolution, 15(5), e71378. 10.1002/ece3.71378

Reichard, J. D., Fellows, S. R., Frank, A. J., & Kunz, T. H. (2010). Thermoregulation during Flight: Body Temperature and Sensible Heat Transfer in Free-Ranging Brazilian Free-Tailed Bats (Tadarida brasiliensis). Physiological and Biochemical Zoology, 83(6), 885–897. 10.1086/657253

Salmon, A. B., Leonard, S., Masamsetti, V., Pierce, A., Podlutsky, A. J., Podlutskaya, N., Richardson, A., Austad, S. N., & Chaudhuri, A. R. (2009). The long lifespan of two bat species is correlated with resistance to protein oxidation and enhanced protein homeostasis. The FASEB Journal, 23(7), 2317–2326. 10.1096/fj.08-122523

Santillán, D. D. M., Lama, T. M., Guerrero, Y. T. G., Brown, A. M., Donat, P., Zhao, H., Rossiter, S. J., Yohe, L. R., Potter, J. H., Teeling, E. C., Vernes, S. C., Davies, K. T. J., Myers, E., Hughes, G. M., Huang, Z., Hoffmann, F., Corthals, A. P., Ray, D. A., & Dávalos, L. M. (2021). Large-scale genome sampling reveals unique immunity and metabolic adaptations in bats. Molecular Ecology, 30(23), 6449–6467. 10.1111/mec.16027

Scheben, A., Mendivil Ramos, O., Kramer, M., Goodwin, S., Oppenheim, S., Becker, D. J., Schatz, M. C., Simmons, N. B., Siepel, A., & McCombie, W. R. (2023). Long-Read Sequencing Reveals Rapid Evolution of Immunity- and Cancer-Related Genes in Bats. Genome Biology and Evolution, 15(9), evad148. 10.1093/gbe/evad148

Schlee, M., & Hartmann, G. (2016). Discriminating self from non-self in nucleic acid sensing. Nature Reviews Immunology, 16(9), 566–580. 10.1038/nri.2016.78

Seltmann, A., Troxell, S. A., Schad, J., Fritze, M., Bailey, L. D., Voigt, C. C., & Czirják, G. Á. (2022). Differences in acute phase response to bacterial, fungal and viral antigens in greater mouse-eared bats (Myotis myotis). Scientific Reports, 12(1), 15259. 10.1038/s41598-022-18240-6

Skoufos, G., Kakoulidis, P., Tastsoglou, S., Zacharopoulou, E., Kotsira, V., Miliotis, M., Mavromati, G., Grigoriadis, D., Zioga, M., Velli, A., Koutou, I., Karagkouni, D., Stavropoulos, S., Kardaras, F. S., Lifousi, A., Vavalou, E., Ovsepian, A., Skoulakis, A., Tasoulis, S. K., … Hatzigeorgiou, A. G. (2024). TarBase-v9.0 extends experimentally supported miRNA–gene interactions to cell-types and virally encoded miRNAs. Nucleic Acids Research, 52(D1), D304–D310. 10.1093/nar/gkad1071

Snyder-Mackler, N., Burger, J. R., Gaydosh, L., Belsky, D. W., Noppert, G. A., Campos, F. A., Bartolomucci, A., Yang, Y. C., Aiello, A. E., O’Rand, A., Harris, K. M., Shively, C. A., Alberts, S. C., & Tung, J. (2020). Social determinants of health and survival in humans and other animals. Science, 368(6493), Article 6493. 10.1126/science.aax9553

Snyder-Mackler, N., Sanz, J., Kohn, J. N., Brinkworth, J. F., Morrow, S., Shaver, A. O., Grenier, J.-C., Pique-Regi, R., Johnson, Z. P., Wilson, M. E., Barreiro, L. B., & Tung, J. (2016). Social status alters immune regulation and response to infection in macaques. Science, 354(6315), 1041–1045. 10.1126/science.aah3580

Staerk, J., Conde, D. A., Tidière, M., Lemaître, J.-F., Liker, A., Vági, B., Pavard, S., Giraudeau, M., Smeele, S. Q., Vincze, O., Ronget, V., da Silva, R., Pereboom, Z., Bertelsen, M. F., Gaillard, J.-M., Székely, T., & Colchero, F. (2025). Sexual selection drives sex difference in adult life expectancy across mammals and birds. Science Advances, 11(40), eady8433. 10.1126/sciadv.ady8433

Swanepoel, R., Leman, P. A., Burt, F. J., Zachariades, N. A., Braack, L. E. O., Ksiazek, T. G., Rollin, P. E., Zaki, S. R., & Peters, C. J. (1996). Experimental Inoculation of Plants and Animals with Ebola Virus. Emerging Infectious Disease Journal, 2(4), 321. 10.3201/eid0204.960407

Tastsoglou, S., Skoufos, G., Miliotis, M., Karagkouni, D., Koutsoukos, I., Karavangeli, A., Kardaras, F. S., & Hatzigeorgiou, A. G. (2023). DIANA-miRPath v4.0: Expanding target-based miRNA functional analysis in cell-type and tissue contexts. Nucleic Acids Research, 51(W1), W154–W159. 10.1093/nar/gkad431

Thomas, S. P. (1975). Metabolism during flight in two species of bats, Phyllostomus hastatus and Pteropus gouldii. Journal of Experimental Biology, 63(1), 273–293. 10.1242/jeb.63.1.273

Toshkova, N., Zhelyzkova, V., Reyes-Ruiz, A., Haerens, E., de Castro Deus, M., Lacombe, R. V., Lecerf, M., Gonzalez, G., Jouvenet, N., Planchais, C., & Dimitrov, J. D. (2024). Temperature sensitivity of bat antibodies links metabolic state of bats with antigen-recognition diversity. Nature Communications, 15(1), 5878. 10.1038/s41467-024-50316-x

Wilkinson, G. S. (2025). Do greater spear-nosed bats have societies? Animal Behaviour, 229, 123343. 10.1016/j.anbehav.2025.123343

Wilkinson, G. S., & Adams, D. M. (2019). Recurrent evolution of extreme longevity in bats. Biology Letters, 15, 20180860.

Wilkinson, G. S., Adams, D. M., Haghani, A., Lu, A. T., Zoller, J., Breeze, C. E., Arnold, B. D., Ball, H. C., Carter, G. G., Cooper, L. N., Dechmann, D. K. N., Devanna, P., Fasel, N. J., Galazyuk, A. V., Günther, L., Hurme, E., Jones, G., Knörnschild, M., Lattenkamp, E. Z., … Horvath, S. (2021). DNA methylation predicts age and provides insight into exceptional longevity of bats. Nature Communications, 12(1), Article 1. 10.1038/s41467-021-21900-2

Wilkinson, G. S., Adams, D. M., & Rayner, J. G. (2024). Sex, season, age and status influence urinary steroid hormone profiles in an extremely polygynous neotropical bat. Hormones and Behavior, 164, 105606. 10.1016/j.yhbeh.2024.105606

Wilkinson, G. S., & South, J. M. (2002). Life history, ecology and longevity in bats. Aging Cell, 1(2), 124–131. 10.1046/j.1474-9728.2002.00020.x

Xie, J., Li, Y., Shen, X., Goh, G., Zhu, Y., Cui, J., Wang, L.-F., Shi, Z.-L., & Zhou, P. (2018). Dampened STING-Dependent Interferon Activation in Bats. Cell Host & Microbe, 23(3), 297–301.e4. 10.1016/j.chom.2018.01.006

Xu, S., Hu, E., Cai, Y., Xie, Z., Luo, X., Zhan, L., Tang, W., Wang, Q., Liu, B., Wang, R., Xie, W., Wu, T., Xie, L., & Yu, G. (2024). Using clusterProfiler to characterize multiomics data. Nature Protocols, 19(11), 3292–3320. 10.1038/s41596-024-01020-z

Zhang, G., Cowled, C., Shi, Z., Huang, Z., Bishop-Lilly, K. A., Fang, X., Wynne, J. W., Xiong, Z., Baker, M. L., Zhao, W., Tachedjian, M., Zhu, Y., Zhou, P., Jiang, X., Ng, J., Yang, L., Wu, L., Xiao, J., Feng, Y., … Wang, J. (2013). Comparative Analysis of Bat Genomes Provides Insight into the Evolution of Flight and Immunity. Science, 339(6118), 456–460. 10.1126/science.1230835

## Supporting References

Giles, K. M., Brown, R. A., Ganda, C., Podgorny, M. J., Candy, P. A., Wintle, L. C., Richardson, K. L., Kalinowski, F. C., Stuart, L. M., Epis, M. R., Haass, N. K., Herlyn, M., & Leedman, P. J. (2016). microRNA-7-5p inhibits melanoma cell proliferation and metastasis by suppressing RelA/NF-κB. Oncotarget, 7(22), 31663–31680. 10.18632/oncotarget.9421

Kacprzyk, J., Hughes, G. M., Palsson-McDermott, E. M., Quinn, S. R., Puechmaille, S. J., O’Neill, L. A. J., & Teeling, E. C. (2017). A Potent Anti-Inflammatory Response in Bat Macrophages May Be Linked to Extended Longevity and Viral Tolerance. Acta Chiropterologica, 19(2), 219–228. 10.3161/15081109ACC2017.19.2.001

